# Molecular and Structural Basis of Cardiac Remodelling in Niemann-Pick Type C

**DOI:** 10.64898/2026.07.05.736597

**Authors:** Qianqian Song, Ishika Prachee, Karolina M Stepien, Neil Herring, Alfonso Bueno-Orovio, Rebecca A Capel, David Priestman, Thamali Ayagama, Laura Bell, Victoria S. Rashbrook, Reuben Bush, Duncan B. Sparrow, Claire Smith, Dave Smith, Emily Akerman, Jianshu Hu, Charalampos Sigalas, Reena Sharma, Peter Woolfson, Ming Lei, Frances M Platt, Rebecca AB Burton

## Abstract

**Key Point Summary:** Niemann-Pick disease type C (NPC) patients showed a high prevalence of ECG abnormalities, with additional echocardiographic evidence of altered left-ventricular structure and function.

*Npc1^-/-^* mouse hearts exhibited age-associated glycosphingolipid accumulation accompanied by marked myocardial fibrosis and increased collagen deposition.

*Ex vivo* electrophysiology revealed QT prolongation and atrioventricular conduction defects in *Npc1^-/-^* hearts, particularly under β-adrenergic stress.

Transcriptomic profiling identified inflammatory and fibrotic pathway activation consistent with the structural and electrophysiological abnormalities observed.

Together, these findings demonstrate previously unrecognised cardiac involvement in NPC and support routine cardiac screening to improve clinical management.

Niemann-Pick disease type C (NPC) is a rare autosomal recessive neurodegenerative lysosomal storage disease caused by pathogenic variants in *NPC1* or *NPC2*. Sudden death can occur due to seizures, but cardiac involvement has not been well defined.

We performed 12-lead electrocardiograms (ECG) in 14 adult NPC patients (8 male, 6 female). Cardiac structure and function were examined in *Npc1^-/-^* adult mouse hearts, alongside wild-type controls. Glycosphingolipid accumulation was quantified by high-performance liquid chromatography, fibrosis and collagen deposition were quantified using Masson’s Trichrome (M&T) and Picrosirius Red (PR) staining. Whole-heart morphology, including chamber size and wall thickness, was assessed. *Ex vivo* ECG recordings assessed conduction abnormalities and arrhythmias. RNA-seq transcriptomics characterised molecular pathways altered in *Npc1^-/-^* hearts.

8/14 patients showed ECG abnormalities including abnormal QRS transitions (N=8), increased QRS amplitude (N=4), fascicular block (N=2), and abnormal T wave inversion (N=1). 13 patients also had transthoracic echocardiograms identifying mildly impaired LV systolic function (N=2) and increased wall thickness/LV mass (N=4). In *Npc1^-/-^* mice, age-related glycosphingolipid accumulation was associated with pronounced ventricular fibrotic remodelling. There was a significant increase in stained connective tissue area and connective tissue to cardiac tissue ratio in both M&T and PR staining. ECG from Langendorff-perfused *Npc1^-/-^* hearts showed QT prolongation and atrioventricular conduction abnormalities under isoprenaline stress. Transcriptomics revealed major changes in *Npc1^-/-^* hearts, consistent with histological fibrosis and linking NPC to inflammation-driven remodelling and arrhythmogenesis.

These findings support routine cardiac screening in NPC patients and highlight the need for further studies to improve management and treatment.

## Introduction

Lysosomal storage disorders (LSD) are a group of inherited metabolic diseases characterised by the accumulation of macromolecules within lysosomes, disrupting normal lysosomal homeostatic functions (Platt *et al*., 2018). Cardiac abnormalities occur in several LSD including sphingolipid storage disorders such as Anderson-Fabry disease, Gaucher’s disease and Acid Spingomyelinase deficiency (ASMD, known as Niemann-Pick diseas type B) (Veinot & Nair, 2020). Anderson-Fabry disease is an X-linked LSD resulting from reduced α-galactosidase A activity, an enzyme that catabolises globotriasylceramide (Gb3) (Tuttolomondo *et al*., 2021). This leads to the accumulation of Gb3 in the lysosome. Its cardiac manifestations include left ventricular hypertrophy (LVH), heart failure and arrhythmia (Pieroni *et al*., 2021; Tuttolomondo *et al*., 2021). Critically, LVH and heart failure are the leading cause of death in Fabry patients (Pieroni *et al*., 2021; Tuttolomondo *et al*., 2021). Gaucher’s disease, caused by mutations in the *GBA* gene, leads to reduced β-glucocerebrosidase activity; this results in the accumulation of glucosylceramide within macrophages. Cardiovascular Gaucher disease (CGD), caused by a D409H (p.Asp409His) variant, is the rarest type of Gaucher disease. CGD involves calcification and fibrosis of the aortic and mitral valves, and pulmonary hypertension (Abrahamov *et al*., 1995; Kurolap *et al*., 2019). ASMD is a rare progressive genetic disorder, caused by a deficiency of acid sphingomyelinase. Some of these patients have cardiac diseases including coronary artery disease, with reports that cardiac disease accounts for >7 % of deaths among adults with chronic visceral or chronic neurovisceral ASMD (McGovern *et al*., 2017).

Niemann-Pick disease type C (NPC) is an autosomal recessive LSD associated with pathogenic variants in two genes: *NPC1* (95 % of cases), which encodes the late endosomal/lysosomal transmembrane protein, and *NPC2*, which encodes a soluble protein with cholesterol binding properties (Vanier *et al*., 1996; Sleat *et al*., 2004). The exact function of these proteins remains incompletely understood but it is apparent they are required for the removal of lipids from the lysosome to the endoplasmic reticulum (ER). In their absence, multiple lipids accumulate in the lysosome including cholesterol and sphingolipids (Loannou, 2000; Sleat *et al*., 2004; Höglinger *et al*., 2019). Neurological manifestations and neurocognitive decline are the commonest and main complications of the disease.

Fibrotic remodeling of the liver is an established manifestation in NPC patients. Paediatric case records have noted fibrotic changes in liver biopsy specimens (Kelly *et al*., 1993). The records showed that 12 out of 15 children that had persistent liver disease had hepatic fibrosis, with five progressing to cirrhosis (Kelly *et al*., 1993). Another study in France demonstrated similar findings in adult patients; liver biopsy studies were performed in 16 patients, and all the samples demonstrated significant fibrosis (Gardin *et al*., 2023). Furthermore, specific knockdown of NPC1 protein levels in mice in the liver caused hepatic fibrosis, detected by increased collagen deposition on Masson’s trichrome stained tissue sections (Sayre *et al*., 2010). However, cardiac outcomes and cardiac fibrosis have not been examined in NPC patients. Investigation into this is pertinent, as myocardial fibrosis could predispose patients to arrhythmias and heart failure (Hinderer & Schenke-Layland, 2019).

As well as fibrosis, inflammation has been noted as a manifestation of NPC (Platt *et al*., 2016). The activation of the innate immune system is a common feature of LSD (Vitner *et al*., 2010). This has mainly been investigated in the central nervous system as neurodegeneration is a key clinical sign. In a mouse model of NPC, changes in pro-inflammatory gene networks were observed in the cerebellum where extensive neurodegeneration takes place (Cologna *et al*., 2012). Here, a series of detoxification enzymes thought to be associated with oxidative stress were all differentially expressed, including the α family of glutathione S-transferase, which was reduced in *Npc1^-/-^* mice compared to WT mice (Cologna *et al*., 2012). A complementary study examined brain tissue from post-mortem NPC patients and *Npc1^-/-^* mice; a gene associated with inflammation – complement 3 was increased in both (Cologna *et al*., 2014). Anti-inflammatory treatment, such as with non-steroidal medications, delays symptom onset in *Npc1^-/-^* mice (Smith *et al*., 2009), further supporting an inflammatory component to the disease pathogenesis of NPC.

Although fibrosis and inflammation are important components of NPC pathogenesis, they do not provide an explanation for all of the disease features. Abnormal cellular Ca^2+^ signaling is thought to contribute to LSD (Lloyd-Evans & Platt, 2011), as well as various other human diseases, including cardiac issues like arrhythmias and heart failure (Landstrom *et al*., 2017). In cardiomyocytes, lysosomes serve as an intracellular calcium (Ca^2+^) store in addition to the sarcoplasmic reticulum. The second messenger nicotinic acid adenine dinucleotide phosphate (NAADP) triggers Ca^2+^ release from endo-lysosomes via two-pore channels (TPCs). This localised Ca^2+^ signal can then contribute to excitation-contraction coupling by modulating calcium release from the sarcoplasmic reticulum (Macgregor *et al*., 2007; Patel & Cai, 2015). NPC is the first identified human disease linked to impaired lysosomal Ca^2+^ uptake and disrupted NAADP-dependent lysosomal Ca^2+^ release (Lloyd-Evans & Platt, 2011).

A Ca^2+^ defect in NPC disease was first proposed by Lloyd-Evans et al. ^(^Lloyd-Evans *et al*., 2008), who specifically highlighted alterations in lysosomal Ca^2+^ levels. Studies in NPC1-mutant human fibroblasts have shown reduced Ca^2+^ levels in the lysosome; 550 μM in WT and 209 μM in the NPC-1 fibroblasts (Lloyd-Evans *et al*., 2008). Alteration of the NPC1 protein function to create a pharmacologically induced NPC model in macrophages led to a ∼55-60 % reduction in intraluminal Ca^2+^ in the acidic compartment, without a change in pH. Sphingosine storage is responsible for this Ca^2+^ deficit; when mouse macrophages were treated with exogenously added lipids before releasing Ca^2+^, only sphingosine was able to cause the reduced Ca^2+^ phenotype (Lloyd-Evans *et al*., 2008).

As the co-ordination of lysosomal Ca^2+^ signaling is disrupted in NPC and is a regulator of cardiac function (Bers, 2002; Lloyd-Evans *et al*., 2008; Capel *et al*., 2015), we hypothesised that cardiac dysfunction may also occur in NPC.

## Methods

### Resource availability

#### Lead Contacts

Further information and requests for resources and reagents should be directed to and will be fulfilled by the lead contact, Rebecca-Ann B Burton

### Material availability

This study did not generate unique new reagents.

### Data and code availability

GEO # GSE279503; Whole tissue transcriptomic data are available upon request

Fibrotic and morphological remodeling – code for analysis. [https://figshare.com/s/552a96811a6cc3991d1f]

### Study Participants

Fourteen patients (8 male, 6 female) diagnosed with NPC under the care of the adult inherited metabolic diseases department in Salford (UK) underwent 12-lead electrocardiogram (ECG) and transthoracic echocardiography as part of a registered health improvement project at Salford Royal Hospital (Registration number: 25HIP12, See Table 1 which also includes age and sex information). All genotype and therapy information is provided in Appendix Table 1.

**Table 1.** 12-lead electrocardiogram data in NPC patients. Summary table of key electrocardiographic parameters measured for each 14 NPC patients. PR interval (PR, msec), QRS duration (QRSd, msec), QRS axis, the presence of left ventricular hypertrophy (LVH) by voltage criteria, timing V1–6 QRS transition, corrected QT interval (QTc, msec) and the presence of T wave abnormality. Abnormalities are highlighted in bold italics. LVH: left ventricular hypertrophy. TWI: T wave inversion. *Left anterior fascicular block. **Left posterior fascicular block.

| Patient | Age | Gender | PR (msec) | QRSd (msec) | QRS Axis | LVH by voltage | V1-6 QRS transition | QTc (msec) | T wave abnormality |
| --- | --- | --- | --- | --- | --- | --- | --- | --- | --- |
| 1 | 33 | M | 151 | 96 | -120 | Yes | Late* | 443 | No |
| 2 | 36 | M | 152 | 117 | -14 | Yes | Late | 428 | No |
| 3 | 39 | M | 143 | 103 | 43 | No | Normal | 401 | No |
| 4 | 33 | F | 130 | 96 | 79 | No | Late | 403 | No |
| 5 | 18 | M | 142 | 77 | 61 | No | Early | 421 | TWI V1-3 |
| 6 | 57 | M | 155 | 81 | 34 | No | Normal | 411 | No |
| 7 | 40 | F | 138 | 94 | 78 | No | Normal | 428 | No |
| 8 | 37 | M | 152 | 99 | 101 | Yes | Late** | 409 | No |
| 9 | 50 | F | 148 | 88 | 58 | No | Normal | 424 | No |
| 10 | 19 | M | 144 | 102 | 38 | No | Early | 442 | No |
| 11 | 37 | M | 177 | 104 | -19 | Yes | Early | 376 | No |
| 12 | 41 | F | 101 | 108 | 31 | No | Normal | 455 | No |
| 13 | 53 | F | 161 | 100 | 26 | No | Normal | 394 | No |
| 14 | 36 | F | 146 | 92 | 19 | No | Late | 392 | No |

### Animal model

All animal studies were conducted using protocols approved by the UK Home Office for the conduct of regulated procedures under licence (Animal scientific Procedures Act, 1986). Animals were humanely killed via cervical dislocation, a Schedule 1 method as defined under the UK Animals (Scientific Procedures) Act 1986. All experimental protocols were approved by the University of Oxford, Procedures Establishment License (PEL) Number XEC303F12. Homozygous *Npc1^-/-^* mice used in these studies were bred from Heterozygous BALB/cNctr-*Npc1^m1N^*/J mice (termed *Npc1^−/−^*mice, also known as *Npc1*^nih^mice; RRID: IMSR_JAX:003092). Mice were bred and housed in individually ventilated cages (IVCs; Thoren, Hazleton, PA, USA) under non-sterile conditions containing Bcell8 bedding (Anibed, France) and given *ad libitum* access to food (i.e., standard chow) and water. The animals were maintained on a 12:12 light: dark cycle. Mice used in electrophysiology studies were at ages of 3, 7, and 9 weeks. For lipids quantification, samples were taken from both male (N=3) and female (N=3) mice; all other experiments in this study used male mice.

## Methods Details

### High-performance Liquid Chromatography Quantification of Glycosphingolipids

Glycosphingolipids (GSLs) were extracted from 7-week-old mouse hearts and analysed using Normal-Phase High-Performance Liquid Chromatography (NP-HPLC) following published methods (Neville *et al*., 2004). Whole heart tissue was homogenized in deionised water and GSls extracted by the addition four times volume of a 1:2 ratio chloroform/methanol (C/M) solution. This was kept overnight at 4°C. The extracts of the mixture were then centrifuged to remove precipitated protein. 0.5 mL of PBS and 0.5 mL of chloroform were added to the supernatant followed by a 3000-rpm centrifugation. The lower phase was carefully removed and dried under a stream of oxygen-free nitrogen in a heating block (42°C). Once dry, lower-phase GSLs were resuspended in 20 μL 1:3 ratioed C/M, and combined with the upper phase. Following this, glycosphingolipids were isolated using C18 columns (Telos, Kinesis, UK). The mixed phase (lower/upper) was loaded onto a column. Bounded GSLs were then extracted from C18 columns by adding 2×1 mL of 98:2 ratio of C/M, 2×1 mL of 1:3 ratio of C/M, and 1 mL of pure methanol. The eluates were dried down under nitrogen and digested with recombinant Endoglycoceramidase I (rEGCase I) (GenScript, Rijswijk, NL) in 90 µL of digestion mix (0.47 mg/mL rEGCase I, 50 mM sodium acetate pH 5.2, and 0.6 % Triton X-100) and incubated at 37°C for 16 hours. The released glycans were then 2AA-labelled (30 mg/mL anthranilic acid (2AA) and 45 mg/mL sodium cyanoborohydride) dissolved in labelling buffer (4 % sodium acetate, 2 % boric acid in methanol) and heated for one hour at 80°C. The labelling mix was allowed to cool to room temperature and mixed with 3 mL 97:3 ratioed A/W and added to a Discovery DPA-6S-SPE tube (Supelco, PA, USA). The columns were cleaned with 4 mL 95:5 ratio of A/W to remove free 2AA. Labelled glycans were eluted from the column with 0.6 mL of water. We took 60 μL from 0.6 mL sample extraction and added 140 μL acetonitrile to reach an A/W ratio of 70:30. The samples were loaded onto a TSKgel® Column, (Tosoh Bioscience, Griesheim, DE), Waters Alliance 2695 separations module and multi-fluorescence detector set at Ex 360/Em 425 nm to separate the labelled glycan headgroups liberated from the GSLs following the digest. To calculate molar quantities from peaks in the chromatogram, a calibration standard containing 2.5 pmol 2AA-labelled chitotriose (Ludger, Oxford, UK) for each NP-HPLC run was included. The chromatographic data was processed using Waters Empower software 3 (Waters, Milford, MA, USA).

### Histological Analysis of the *Npc1^-/-^* Ventricles

Hearts from 7 weeks *Npc1^-/-^* mice were fixed in 4 % paraformaldehyde (PFA) solution and embedded onto a paraffin block. The hearts were serially sectioned (10 μm thickness) and every section was collected on positively-coated slides. Serial sections were either stained with Trichrome Stain Kit, Masson-Light Green (RBK-0601-00B, Cell Path) or in Picrosirius Red solution (Abcam, ab246832) to identify collagen. With Mason’s Trichrome, the stain colour is blue-green for collagen, pink for myocytes, orange for cytoplasm (highlighting nonmyocytes), and blue-black for nuclei. With picrosirius red staining, in bright field microscopy, collagen is red on a pale-yellow background (Figure 2). Some histological images lacked embedded scale bars, necessitating manual estimation of measurements using calibrated image analysis software (e.g., ImageJ, NDP.view2 Viewing software U12388-01). These estimations relied on available image data from source slides, consistent magnification, and identifiable anatomical landmarks. As a result, scale bars should be regarded as near-value measurements.

All images were processed by a custom-made automatic segmentation program in Matlab (figshare code, DOI: 10.6084/m9.figshare.26927728). In brief, image segmentation was automated in Matlab 2021a, exploiting built-in routines of its image processing toolbox. For each histology section, the region of interest (ROI) was delimited as follows. Firstly, images were greyscale converted, and edges were detected as zero-intensity crossings after filtering with a Laplacian of Gaussian filter. This was followed by image binary dilation, filling, and erosion operations to yield a binary mask of the external envelope of the tissue slice. Similarly, any internal cavities were automatically detected by applying the same principles to the image complement (negative) of the greyscale image, followed by binary erosion and dilation operations. The final ROI was defined as the difference between the external and internal binary masks and applied to each of the RGB channels of the original true-colour image.

For connective tissue segmentation, a decorrelation stretch was first applied to the masked RGB image, followed by adaptive histogram equalisation of its lightness component. These pre-processing steps were performed to account for variable inter-section staining and to enhance the colour differences between different tissue types in thetrichrome tissue histology sections. The R, G, and B components of the resulting colour-enhanced image were used as input features for k-means clustering with pre-specified centroids start. The resulting clusters were consecutively assigned to the following segmented categories: connective tissue, cardiomyocytes, and other cardiac tissue. Statistics of percentual tissue occupancy were computed as the quotient between the numbers of pixels in each of these categories concerning those in the original ROI. All images were processed without down-sampling at their original resolution (3000×3000 pixels).

### Morphological assessment of the *Npc1^-/-^* Hearts

High resolution episcopic microscopy (HREM) was conducted following established protocols (Weninger & Mohun, 2007). Specimens were fixed overnight at room temperature using Bouin’s fixative (HT10132, Merck), thoroughly rinsed in PBS, and then subjected to gradual dehydration in a methanol series. Subsequently, samples were embedded utilizing a JB-4 kit (Catalogue number 00226-1, Polysciences) through an overnight infiltration at 4°C with a 50:50 mixture of methanol and infiltration solution (JB-4 Solution A plus 1.25 % (w/v) Benzoyl Peroxide Plasticizer). Following a 60-minute wash at 4°C in 100 % infiltration solution, the samples were left to incubate for up to 1 week at 4°C in fresh infiltration solution. The embedding process involved the addition of 6 % (v/v) JB-4 solution B to the infiltration solution to initiate polymerization. Analysis was carried out using a 3D Optical high-resolution episcopic microscopy imaging system (Indigo Scientific). HREM data were evaluated using Horos 3.3.6 (https://horosproject.org), and Amira for Life & Biomedical Sciences version 2019.4 (Thermo Fisher Scientific). Example videos of wild type (WT) and *Npc1^-/-^* hearts provided in Supplementary Videos 1 and 2.

### *Ex vivo* Electrocardiogram

Hearts were taken from 7 weeks old WT and *Npc1^-/-^* mice and place on a Langendorff system perfused in Krebs solution (in mM: NaCl 119, NaHCO3 25, KCl 4.0, MgCl2 1.0, CaCl2 1.8, and glucose 10) maintained at pH 7.4 by bubbling with 95 % O_2_-5 % CO_2_. ECG were recorded using bipolar electrogram (BEG) electrodes with 1mm interpole spacings placed atthe right outflow tract. Post cancellation, hearts were left to stabilise for 5-10 minutes before live recording. The resulting electrogram signals were amplified for band-pass filtering between 0.5 Hz and 1 kHz. All signals were recorded using an NL100AK head stage and NL104 amplifier unit, band-pass filtered (Neurolog NL 125/6 Filter, Neurolog, Hertfordshire, UK), and then digitized at 5 kHz by a micro 1401plus MKII laboratory interface (Cambridge Electronic Design, Cambridge, UK) using Spike2 software (Cambridge Electronic Design). Electrocardiogram traces were recorded in Spike2 software with exact time point which were used to calculated RR (heart rate), PR, and QT interval of the hearts.

### Transcriptomic Analysis

#### Sample preparation

Frozen cardiac tissue was collected without the RNase contamination and thawed using an RNAlater^TM^ stabilisation solution (AM7020, Invitrogen). Samples were homogenised using bead disruption, and in 1 mL of Trizol (15596026, Thermo Fischer), approximately 50 – 100 mg of tissue was solubilised. The samples were rested in room temperature for 10 minutes and centrifuged at 12,000 x g for 5 minutes at 4°C. 200 μL chloroform was added to the supernatant and the mixture was centrifuged again at 12,000 g at 4°C for 15 minutes. The upper aqueous phase was separated, 1:1 volume ice-cold isopropanol was added, and gently mixed. This procedure was followed by centrifugation at 12,000 g for 30 minutes at 4°C, and the collected RNA pellet was washed with 75 % ethyl alcohol (10048291, Thermo Fisher) and dried.

#### Qualification and Quantification of mRNA

The isolated RNA was screened for contaminants and degradation levels using 1% agarose. A NanoPhotometer® spectrophotometer (IMPLEN, CA, USA) was used to check the purity. Finally, the RNA Nano 6000 Assay Kit of the Bio-analyser 2100 system (Agilent Technologies, CA, USA) was used to detect the RNA integrity and assess quantification. Sequence libraries were prepared using the NEBNext® Ultra TM RNA Illumina® (NEB, USA) Library Prep Kit. Furthermore, using a PE Cluster Kit cBot-HS, all the samples were clustered and sequenced using an Illumina. Fast’ was used to remove the poly-N, and an adapter reads from the raw data and processes it. The Q score at 20 and 30 were identified as clean. The Genome web browser National Centre for Biotechnology Information/European Molecular Biology Laboratories-European Bioinformatics Institute (NCBI/Ensembl-EBI) was used as the reference genome with the HISAT2 programme (daehwankimlab.github.io/hisat2/ manual/). The gene Gene Ontology and KEGG analysis were performed using g:GOSt on web server g:Profiler (https://biit.cs.ut.ee/gprofiler/gost). Supplementary files S1-S9 related to Figure 4-6, contain all related RNA Seq data.

### RT-qPCR

Extraction and purification of total RNA from whole heart tissue were conducted using RNeasy Plus Mini Kit (Qiagen Inc., USA). RNA concentration was measured before RNA is reverse transcribed to cDNA using a High-Capacity cDNA Reverse Transcription Kit (Applied Biosystems™, UK). cDNA sample were assessed for gene of interest (*Lgals3*) and normalized with house keeping gene *Gapdh* using Polymerase Chain Reaction (rt-PCR) Master Mix TaqMan™ Fast Advanced (Thermo Fischer, UK). Primer used is *Lgals3* (TaqMan, Assay ID: Mm00802901_m1, Thermo Fisher, UK) which is normalized with the reference gene *Gapdh* (TaqMan, Assay ID: Mm99999915_g1, Thermo Fisher, UK) expression. Non-parametric test was performed to asses significance.

## Data Analysis and Statistics

### Patient studies

Patient demographic data is presented as median [interquartile range].

### Animal studies

Each histology slice was quantified and included as n=1, therefore data n number represents the number of slices analyzed per group of interest whilst N=number of animals. For morphology, glycosphingolipid chromatography, ECG and transcriptomic data, samples were extracted from individual animals and quantified, a single mouse represents N=1, with several N’s per study method. In this study, n represents the number of biological replicates where N equals the number of individual animals used. All statistics for the histology, morphology, lipid chromatography and ECG interval calculation of the ECG data were analyzed using two tailed unpaired student t-test assuming equal variance, and ECG abnormal events prevalence were analyzed using Fisher’s exact test. The significance is recognized at *p*<0.05 (*), *p*<0.01(**), *p*<0.001(***), and *p*<0.0001(****). Results with limited number of samples (n or N<5) were analyzed using non-parametric test without assuming followed a normal distribution. The statistical calculations and plotting were done using GraphPad Prism (Version 10.2.2 for Mac), and data are presented as mean±SD (standard deviation).

## Results

### 1. Electrocardiograms and transthoracic echocardiograms in NPC patients

Fourteen adult NPC patients (8 male, 6 female), with a median age of 37 [34-41] years old and a body mass index of 25.6 [20.6-29.1] kg/m^2^, underwent 12-lead ECGs (Table 1). The genotypes of each patients and their treatments are summarised in Appendix Table 1. One patient had an additional cardiomyopathy-related variant (an autosomal dominant variant in the *MYBPC3* gene). All patients presented a stable resting beating rate (73 [68-80] beats per mimute; N=14) and none were hypertensive (blood pressure 117 [110-121] / 76 [73-79] mmHg). Eight out of the fourteen presented a form of ECG abnormality (Table 1). These included abnormal QRS transitions in the chest leads (N=8), LVH by voltage (N=4), anteroseptal T wave inversion, left anterior fascicular block, and left posterior fascicular block (N=1 each). Thirteen of the fourteen patients went on to have transthoracic echocardiograms. Mildly impaired LV systolic function was identified in two patients and increased septal wall thickness in 4 patients, associated with mild and moderately increased LV mass in 2 of these patients (Table 2). There was no significant (>mild) valvular disease.

**Table 2:** Transthoracic echocardiography data in NPC patients. Summary table of key echocardiography parameters measured for each of 13 NPC patients. Left ventricular ejection fraction (LVEF, %), left ventricular internal diameter in diastole (LVIDd, mm), interventricular septal thickness in diastole (IVSd, mm), left ventricular mass index (LVMi, g/m²), and left atrial diameter (LAd, mm). Abnormalities highlighted in bold italics. LVEF: left ventricular ejection fraction. LVIDd: left ventricular internal diameter in end diastole, IVSd: interventricular septal diameter in end diastole. LVMi: left ventricular mass index. LAd: left atrial diameter.

| Patient | LVEF | LVIDd | IVSd | LVMi | LAd |
| --- | --- | --- | --- | --- | --- |
|  | (%) | (mm) | (mm) | (g/m<sup>2</sup>) | (mm) |
| 1 | 55 | 27 | **23** | **139** | 35 |
| 2 | **45** | **60** | **11** | **126** | 3 |
| 3 | 59 | 51 | 11 | 98.4 | 35 |
| 4 | 76 | 46 | 9 | 70 | 29 |
| 5 | 55 | 32 | 7 | 42 | 24 |
| 6 | 57 | 39 | **12** | 59 | 37 |
| 7 | - | - | - | - | - |
| 8 | 55 | 45 | 9 | 68 | 26 |
| 9 | 60 | 37 | **10** | 65 | 18 |
| 10 | **45** | 37 | 8 | 55 | 30 |
| 11 | 55 | 42 | 9 | 58 | 25 |
| 12 | 55 | 47 | 7 | 54 | 29 |
| 13 | 55 | 40 | 9 | 61 | 35 |
| 14 | 55 | 45 | 5 | 53 | 34 |

### 2. Lipid accumulation is observed in the hearts of *Npc1^-/-^* mice

Accumulation of glycosphingolipids has been shown to occur in various organs of NPC patients, including the liver, kidney and brain (Pacheco & Lieberman, 2008). A biochemical lipid analysis, targeting glycosphingolipids, was therefore performed on *Npc1^-/-^* mice cardiac tissue to assess whether this occurs in NPC hearts (Figure 1A). Due to the severity of the genotype, *Npc1^-/-^* mice have a lower lifespan compared to WT mice, living approximately 10-12 weeks. Therefore, for this study we chose mice aged at weeks 3, 7 and 9 for studying the disease progression and phenotype. Glycosphingolipid levels increased from 7 weeks of age (N=6) and continued to accumulate up to 9 weeks of age (N=6) relative to age matched controls (Figure 1B).

**Figure 1.**
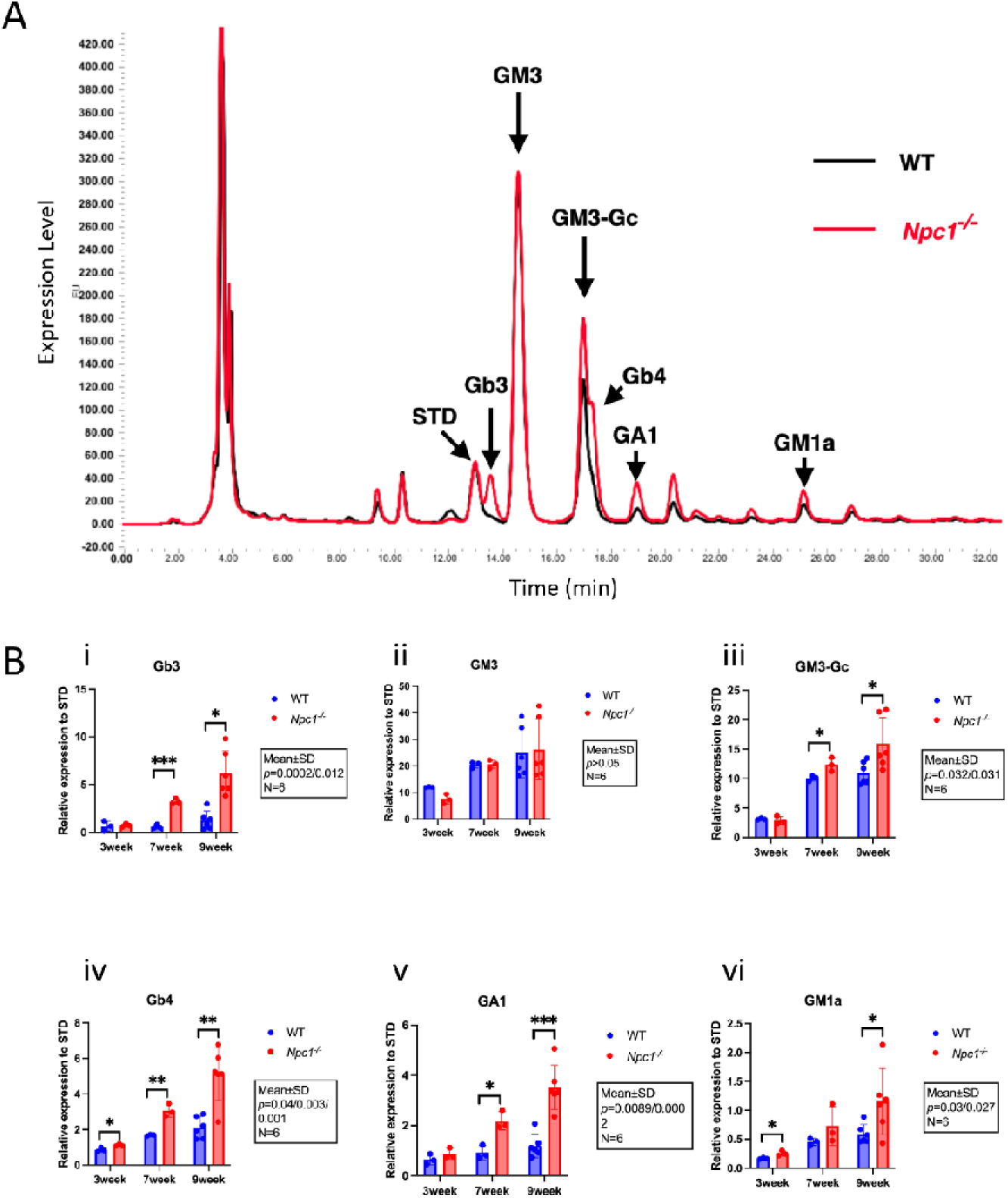
Glycolipids are accumulated in *Npc1^-/-^* hearts. A. Representative high performance liquid chromatography traces from hearts of WT (black) and *Npc1^-/-^* (red) mice. B. Levels of lipids, Gb3 (i), GM3 (ii), GM3-Gc (iii), Gb4 (iv), GA1 (v), GM1a (vi) in WT (N=6, blue) and *Npc1^-/-^* (N=6, red) at 3, 7 and 9 weeks. Data are presented as mean±SD. Student t-test. Expression level of each lipid has been normalized to standard (STD).

In *Npc1^-/-^* hearts, Gb3 (week 7, \*\*\**p*=0.0002; week 9, \**p*=0.012) was the most elevated glycosphingolipid, alongside Gb4 (week 7, \*\**p*=0.003; week 9, \*\**p*=0.001), GA1 (week 7, \*\**p*=0.0089; week 9, *p*=0.0002), GM3-Gc (week 7, \*\*\**p*=0.0002; week 9, \**p*=0.012) and GM1a (week 9, \**p*=0.027) (Figure 1B). Glycosphingolipid analysis presented in Figure 1B are consistent with previous findings in other tissues in NPC patients^(Pacheco & Lieberman, 2008)^. Thus, the *Npc1* gene defect causes altered lipid trafficking in cardiac cells, and has the potential to affect cardiac function at the cellular level. We also performed similar analysis in brain and liver tissue and the results are presented in Figure A1.

### 3. Fibrotic and morphological remodeling in *Npc1^-/-^* mouse hearts

Given the evidence of sphingolipid accumulation in *Npc1^-/-^* cardiac tissue, it is plausible that cellular disruption could lead to pathological cardiac remodeling. To evaluate cardiac morphology, hearts from 7-week-old WT and *Npc1^-/-^* mice were analyzed histologically. WT (N=3) and *Npc1^-/-^* (N=3) hearts, were sectioned, and ventricles were stained with Masson’s Trichrome (M&T) and Picrosirius Red (PR) stains, to assess both myocardial fibrosis and collegen deposition, respectively. High-resolution images were processed by a custom-made automatic separation/detection programme in Matlab (code provided). There was a significant increase in both the percentage area of stained connective tissue, and increased connective tissue to cardiac tissue ratio, in both M&T (\*\*\*\**p*<0.00001, n=116/93 sections, N=3 hearts; Figure 2A) and PR staining (\*\*\*\**p*<0.00001/=0.0003, n=14/18 sections, N=3 hearts; Figure 2B). Both histological analyses suggest that *Npc1^-/-^* ventricular tissue undergoes fibrotic remodeling from 7 weeks of age in mice, corresponding to the early symptomatic phase of the disease. Additional PR staining on 9-week-old hearts, similarlary showed an increase of fibrosis were in *Npc1^-/-^* mice (Figure A2A and B).

**Figure 2.**
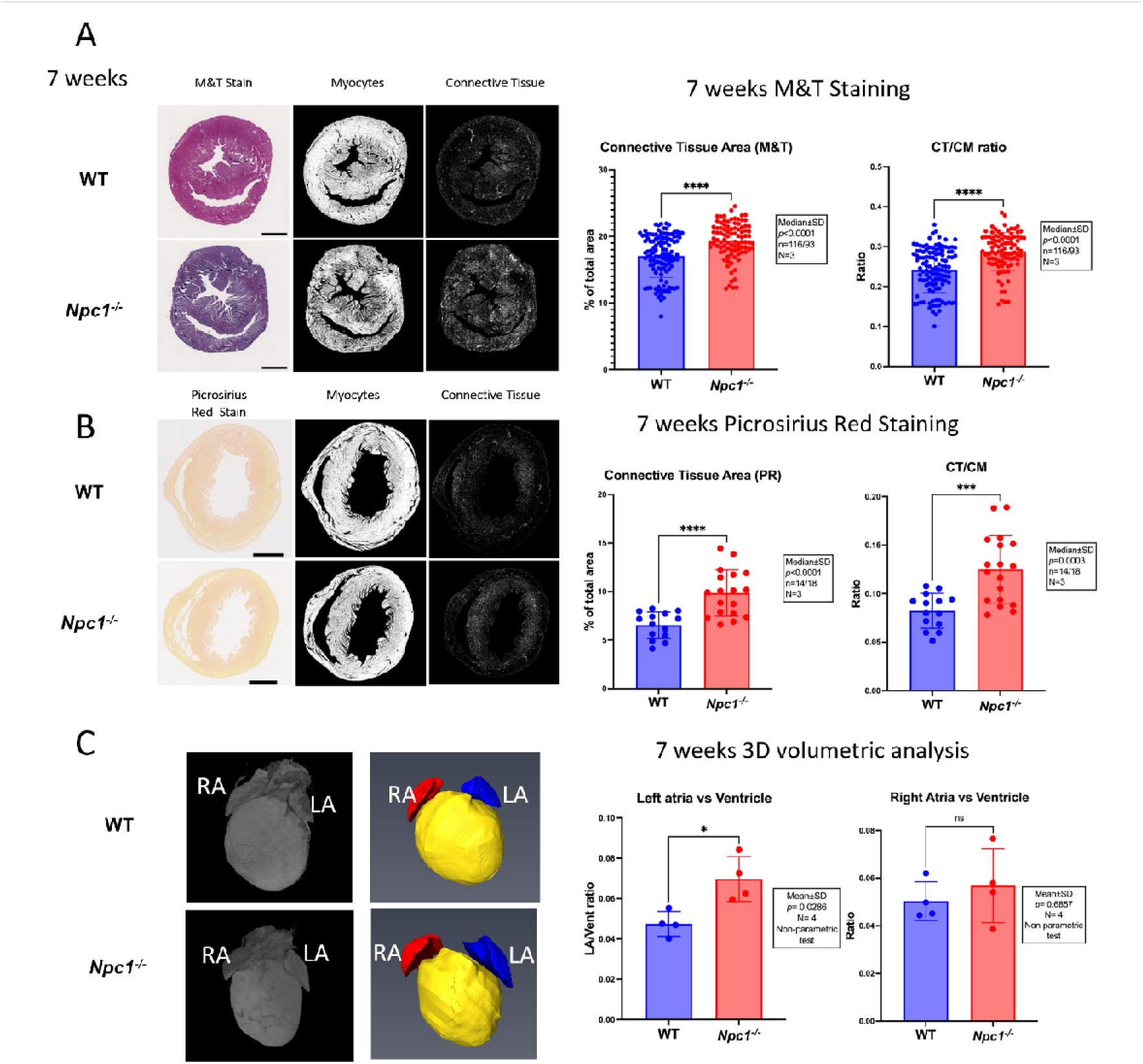
Histology and 3D reconstruction of hearts from *Npc1^-/-^* mice reveal fibrotic and structural remodeling. A. Sections stained with Masson’s Trichrome (M&T) in hearts from 7-week-old WT (N=3, n=116) and *Npc1^-/-^* (N=3, n=93) mice (\*\*\*\**p*<0.0001). B. Sections stained with Picrosirius Red in hearts from 7-week-old WT (N=3, n=14) and *Npc1^-/-^* (N=3, n=18) mice (\*\*\*\**p*<0.0001). C. 3D model generated from High Resolution Episcopic microscopy (HREM) images of WT (N=4) and *Npc1^-/-^* (N=4) hearts using Amira software (\**p*=0.0128 in *Npc1^-/-^* hearts; non-parametric test); LA: left atrium and RA: right atrium. Scale bars in Panel A & B = 1 mm.

### 4. Morphologic assessment through 3D reconstruction using High Resolution Episcopic Microscopy (HREM) reveals gross anatomical changes in *Npc1^-/-^* mouse hearts

To determine if any structural remodeling in atrial tissue is observed, a volumetric analysis through 3D reconstruction was performed (Figure 2C). 7-week-old WT (N=4) and *Npc1^-/-^* (N=4) hearts were analyzed using HREM and 3D models were generated using Amira software. Right and left atrium and ventricle areas were manually segmented for volume comparison. The volume of the ventricle showed no significant difference between WT and *Npc1^-/-^* hearts (Figure A2C, Supplementary Video 1 (WT) and 2 (*Npc1^-/-^*). However, the left atrium to ventricle ratio was significantly larger in the *Npc1^-/-^* group (\**p*=0.0286, N=4, non-parametric t-test). By contrast, the right atria did not show a significant increase in size (Figure 2C). Supporting these data, *Npc1^-/-^* hearts analyzed by both conventional histology or assessed visually also had a visibly enlarged left atrium.

### 5. Abnormal cardiac electrophysiology observed in *Npc1^-/-^* mouse hearts

Fibrotic remodeling and cellular impairment can impact multiple aspects of cardiac function, including the heart’s electrophysiological activity. Cardiac electrophysiology was assessed by conducting *ex vivo* bipolar ECGs on Langendorff-perfused WT and *Npc1^-/-^* hearts (Figure 3A). Recording were obtained from 7-week-old mice in both groups. *Npc1^-/-^* hearts exhibited similar heart rates (HR) to WT hearts (WT=317.1±68.36 bpm, N=10; *Npc1^-/-^* =302.3±63.85, N=9; *p*>0.05), both at baseline and following isoprenaline (Iso, 50 nM) stress (WT=408.9±110.6 bpm, N=10; *Npc1^-/-^* =396.8±98.92, N=9; *p*>0.05) which heart rate had significantly increased (WT vs WT Iso, \*\**p*=0.0033; *Npc1^-/-^* vs *Npc1^-/-^* Iso, \**p*=0.0136, Figure 3B). There was no overall difference in PR interval in either condition in mice with 1:1 AV conduction (Figure 3C; *p*>0.05), however QT interval was also significantly prolonged both at baseline and during Iso stress (Figure 3D; \*\*\*\**p*<0.0001). The incidence of first (>50msec) and second-degree AV blocks was significantly higher following Iso treatment, incease from 2 out of 9 hearts to 5 out of 9 hearts (Figure 3E, \**p*=0.011, Fisher Exact test). ECG trace examples of type of AV blocks observed in *Npc1^-/-^* mice are presented in Figure A3.

**Figure 3.**
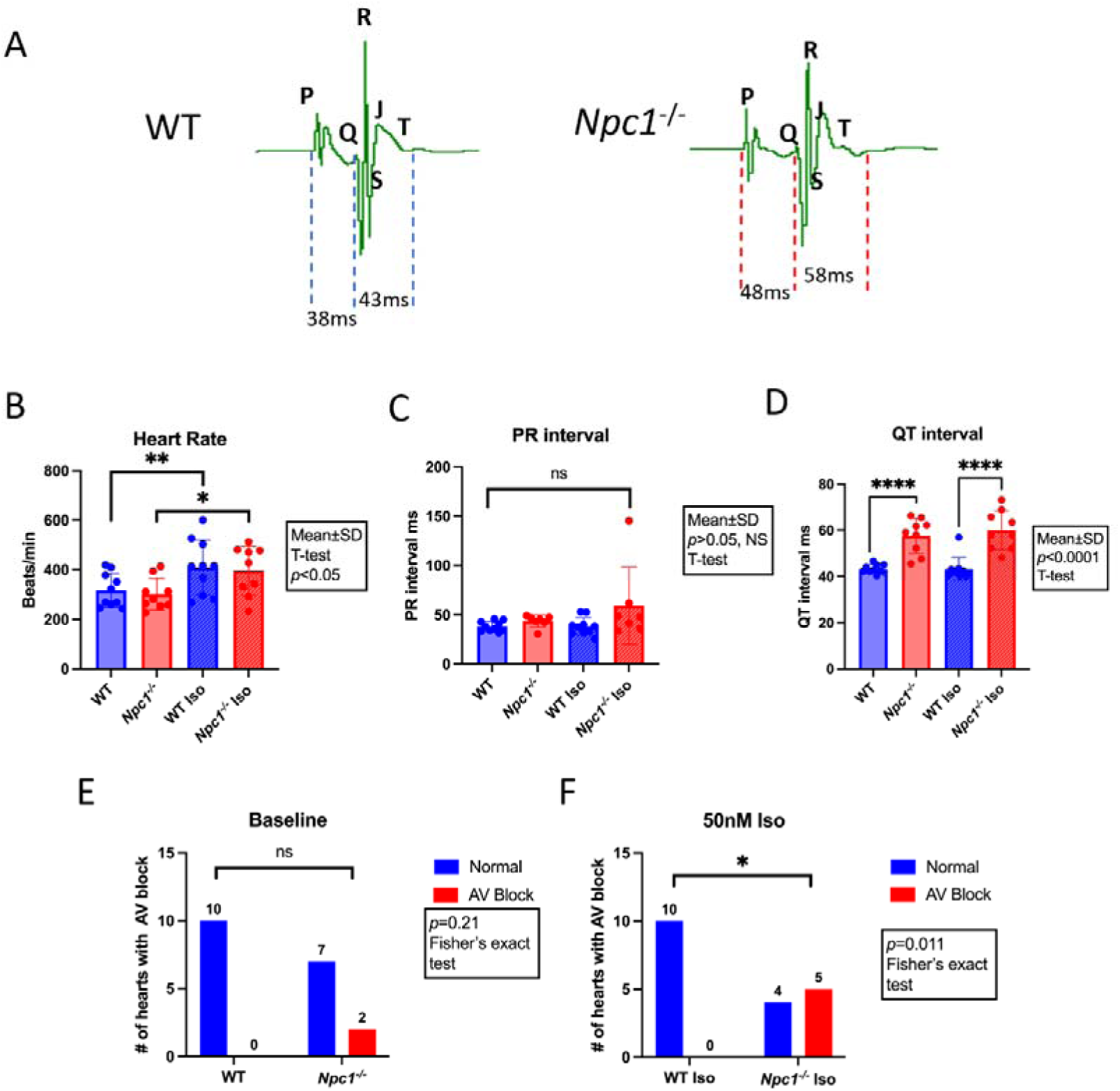
Electrophysiological recordings from *Npc1^-/-^* vs WT hearts. A. Example electrocardiogram trace from WT and *Npc1^-/-^* hearts. B. Heart rate of WT (N=10) and *Npc1^-/-^* (N=9) at baseline and following addition of isoprenaline (Iso, 50 nM; \**p*=0.0136, \*\**p*=0.0033). C. PR intervals in ms of WT (N=10) and *Npc1^-/-^* (N=9) at baseline and following addition of Iso (50 nM). D. QT intervals in ms of WT (N=10) and *Npc1^-/-^* (N=9) at baseline and following addition of Iso (50 nM; \*\*\*\**p<*0.0001). E. Incidence of first- and second-degree AV blocks observed in WT (N=10) and *Npc1^-/-^* (N=9) hearts before Iso. F. Incidence of first- and second-degree AV blocks observed in WT (N=10) and *Npc1^-/-^* (N=9) hearts in the presence of Iso (50 nM);(\**p=*0.011, Fisher exact test).

### 6. Transcriptomic analysis of *Npc1^-/-^* hearts

Morphological and functional experiments presented above demonstrated that *Npc1^-/-^* hearts undergo pathological, morphological and electrophysiological remodeling. To explore the potential pathways leading to this remodeling, transcriptomic profiling was conducted on WT and *Npc1^-/-^* mouse hearts at 3, 7, and 9 weeks of age (N=5 for each age point) representing pre-symptomatic, early symptomatic and late symptomatic stages of disease respectively. Over 5,300 differentially expressed genes (DEGs) were detected, comprising 3,240 DEGs at 3 weeks old, 2,190 DEGs at 7 weeks old, and 5,300 DEGs at 9 weeks old (Figure 4A, Supplementary File 1-3, Figure A4). Only DEGs with padj< 0.05 (-log padj > 1.301) are considered significant and included in pathway analysis. Principal component analysis (PCA) was done using these gene profiles to quantify the differences in sample composition (Figure 4B). Analysis revealed that *Npc1^-/-^* hearts exhibit distinct profiles compared to WT hearts. As showed in the PCA plot, after week 7, all *Npc1^-/-^* samples are distributed at different spacial area on the axis, distant from WT samples, and the distance increased with disease progression. Distribution of up and down-regulated DEGs are showed as a volcano plot, top up/down regulated DEGs are labelled (Figure 4C).

**Figure 4.**
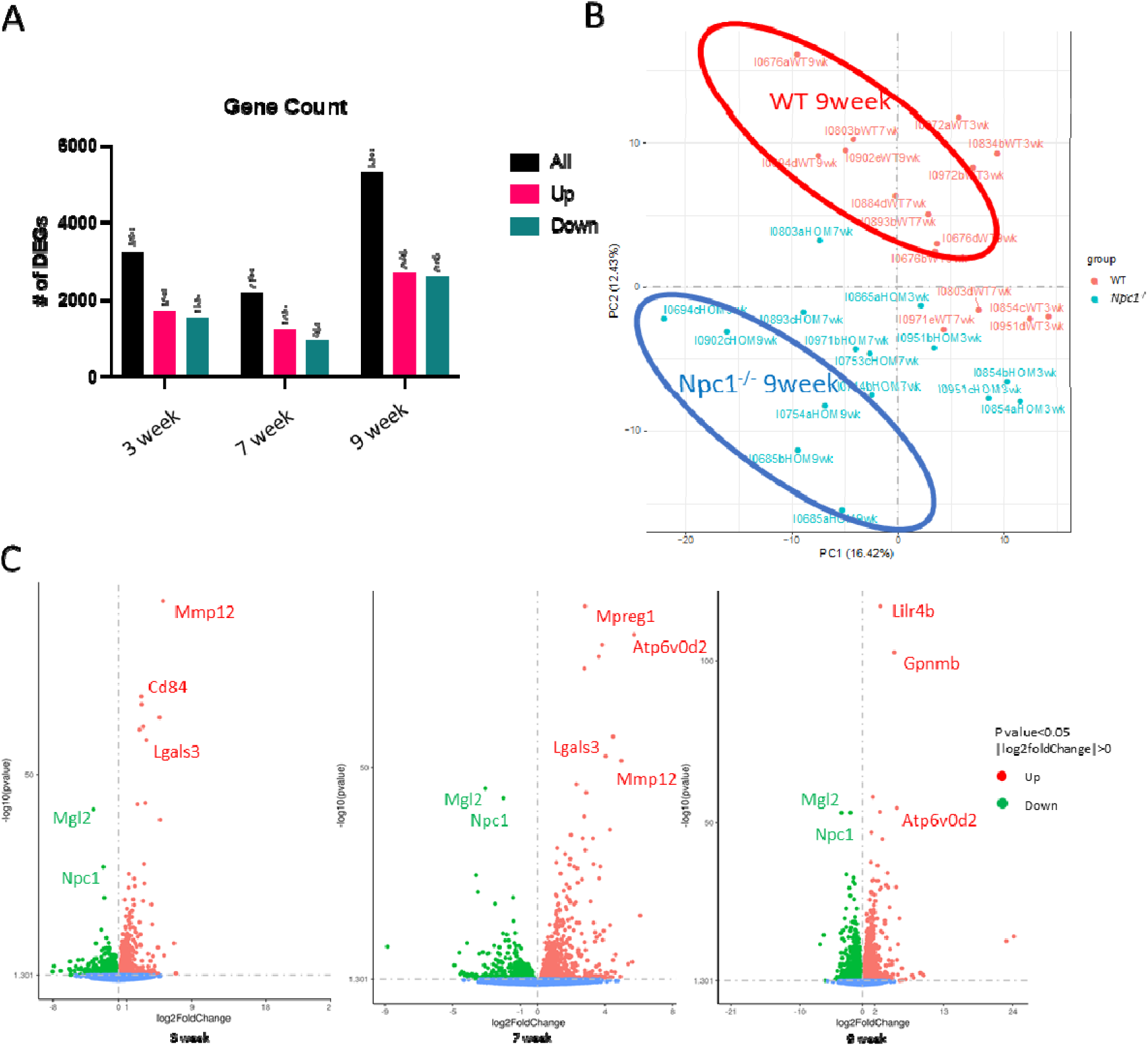
Transcriptomic analysis on *Npc1^-/-^* and WT hearts at 3, 7, and 9 weeks of age. A. Plot of number of differential expressed genes (DEGs) detected in murine *Npc1^-/-^* compared to WT hearts tissue, at 3, 7 and 9 weeks of age. B. Principal component analysis demonstrating the special resolution of WT *Npc1^-/-^* dataset. Distance between each dot represents their data composition differences. C. Volcano plot analysis of WT and *Npc1^-/-^* DEGs showing significantly upregulated (Red) and downregulated (Green) proteins in the *Npc1^-/-^* (*p*<0.05, N=5). The same top up/down regulated DEGs can be found at 3 age points 3,7 and 9 weeks of age.

Further analysis of the transcriptomic profile of *Npc1^-/-^* hearts was performed using GO Biological Process (GO:BP) enrichment analysis, which revealed that inflammation/immune-related pathways predominated (Figure 5 and Figure A5, Supplementary Files 4-6). Among those top regulated GO:BP pathways, we identified hits related to immune and inflammation regulation (Figure 5A); results indicate that immune activation starts as early as week 3 and continues until the late stage at week 9. Several top-regulated DEGs such as *Lgals3* and *Mmp12* (Figure 5Bi and ii), which have both previously been linked to cardiac inflammation and fibrosis, were observed (Mouton *et al*., 2018b; Suthahar *et al*., 2018b). Mgl2 paired with Cx3r1 (Figure 5B iii and iv) has been implicated in cardiac inflammation activation which leads to mitral valve fibrosis and left atrial enlargement (Meier *et al*., 2018). This phenomenon is also observed in our morphological study where left atria are significant larger in *Npc1^-/-^* mice (this study Figure 2C).In addition, *Ccl3* and *Ccl4* expression (Figure 5Biv-vi) were highly upregulated in *Npc1^-/-^* hearts, suggesting activation of the NFkB pathway, a well-recognized regulator of inflammatory responses (Kim *et al*., 2017; Fiordelisi *et al*., 2019). These data are in agreement with the previously reported strong association between inflammation/immune response in NPC disease (Vitner *et al*., 2010). Expression of *Lgals3* was validated using RT-qPCR in independent mouse samples and showed a significant upregulation in *Npc1^-/-^* hearts (N=3) compared to WT (N=5, \**p*=0.0357; non-parametric t-test), consistent with transcriptomic data from whole-heart analysis (Figure A7 and Supplementary File 10).

**Figure 5.**
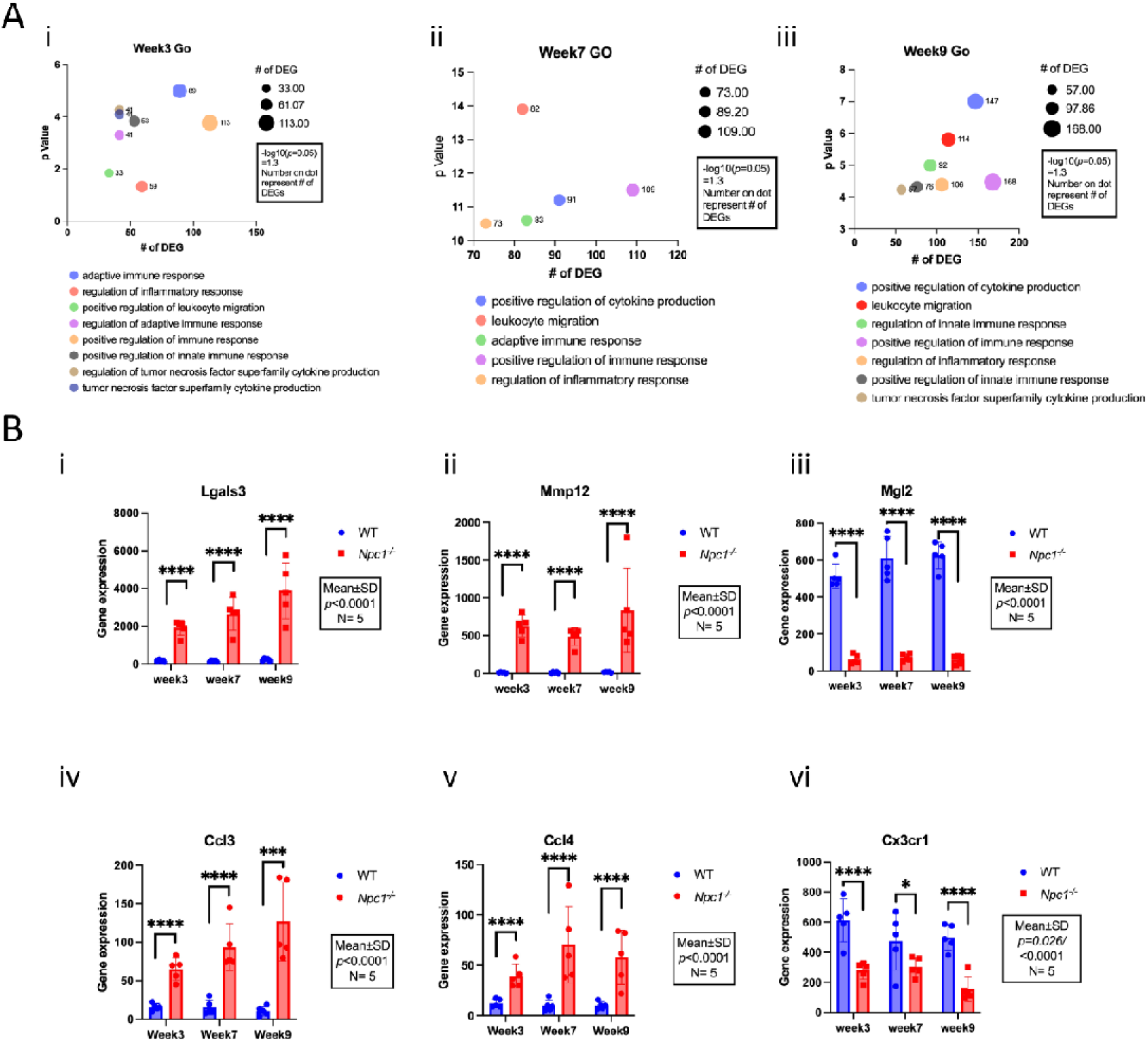
GO-Biological Process Enrichment Analysis on *Npc1^-/-^* and WT hearts transcriptomic. A. Pathway enrichment analysis of DEGs grouped by pathway classification.The number of DEGs associated with each pathway and their corresponding enrichment p-values were analysed at 3 (i), 7 (ii) and 9 (iii) weeks of age. B. Gene expression of *Lgals3* (i)*, Mmp12* (ii), *Mgl2* (iii)*, Ccl3* (iv)*, Ccl4* (v) and *Cx3cr1* (vi) at 3, 7 and 9 weeks of age in WT (N=5) and *Npc1^-/-^* (N=5) mice. Data are presented as mean±SD; Student t-test. *Lgals3*, N=5, *p*<0.0001; *Mmp12*, N=5, \*\*\*\**p*<0.0001; *Mgl2*, N=5, \*\*\*\**p*<0.0001;*Ccl3*, N=5, \*\*\*\**p*<0.0001; *Ccl4*, N=5, \*\*\*\**p*<0.0001; *Cx3cr1*, N=5, \*\*\*\**p*<0.0001.

The relation between cardiac disease and signaling pathways in Npc1^-/-^ mice was further explored using KEGG pathway analysis (Figure 6A and Figure A6). *Atp6v0d2*, a gene encoding a lysosomal proton transporter, showed significant upregulation in *Npc1^-/-^*Hearts (Figure 6Bi). Maintenance of a proton gradient is crucial for maintaining Ca^2+^ concentration in lysosomes (Yang *et al*., 2019; Lloyd-Evans & Waller-Evans, 2020). Hence, alterations in lysosomal proton transport may provide evidence for lysosomal Ca^2+^ dysregulation in NPC diseased cardiomyocytes. From the KEGG analysis, cardiac specific pathways were not activated at week 3, but several DEGs related to cardiac diseases or signaling pathways emerged at week 7 (early symptomatic), including dilated cardiomyopathy (DCM), hypertrophic cardiomyopathy (HCM) and arrhythmogenic right ventricular cardiomyopathy (ARVC). Additionnally, genes *Adcy1* and *Ryr2*, implicated in Cyclic adenosine 3′,5′-monophosphate (cAMP) signaling and Ca^2+^ signaling were detected in and significantly elevated in *Npc1^-/-^* hearts compared to WT at week 9 (Figure 6Bii and iii, Figure A6, Supplementary Files 7-9). In addition to KEGG, at week 9, the analysis revealed pathways significantly involved in extracellular matrix (ECM) organization and TGF-beta activation, both of which are closely related to cardiac fibrosis. This additional bioinformatic analysis of the transcriptomics data provided further evidence that the *Npc1^-/-^* heart undergoes structural and molecular remodeling.

**Figure 6.**
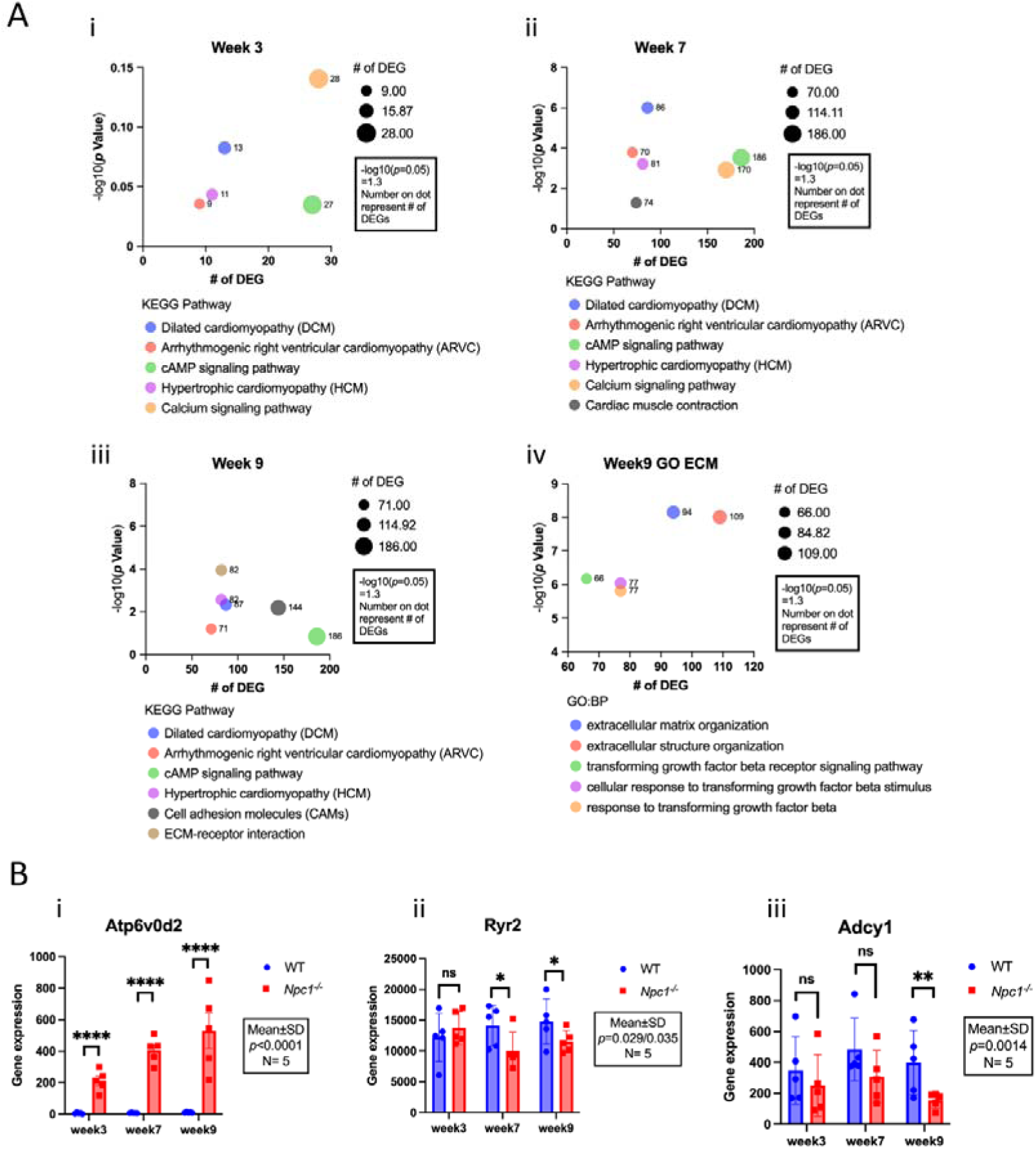
KEGG analysis of cardiac signaling/disease pathways on *Npc1^-/-^* and WT hearts transcriptomic. A. KEGG pathway analysis performed in DEGs from *Npc1^-/-^* mice at 3 (i), 7 (ii), and 9 (iii) weeks of age showing pathways that involve cardiac signaling/disease. From 9 weeks of age, significant involvement of sevel pathways related to extracellular matrix (ECM) remodeling has been found. (iv). B. Gene expression of *Atp6v0d2* (i)*, RyR2* (ii) and *Adcy1* (iii) at 3, 7 and 9 weeks of age WT (N=5) and *Npc1^-/-^* (N=5) mice. Data are presented as mean±SD; Student t-test. *Atp6v0d2*, N=5, *p*<0.0001; *Ryr2*, N=5, \**p*=0.029/0.035; *Adcy1*, N=5, \*\**p*=0.0014;.

**Figure 7.**
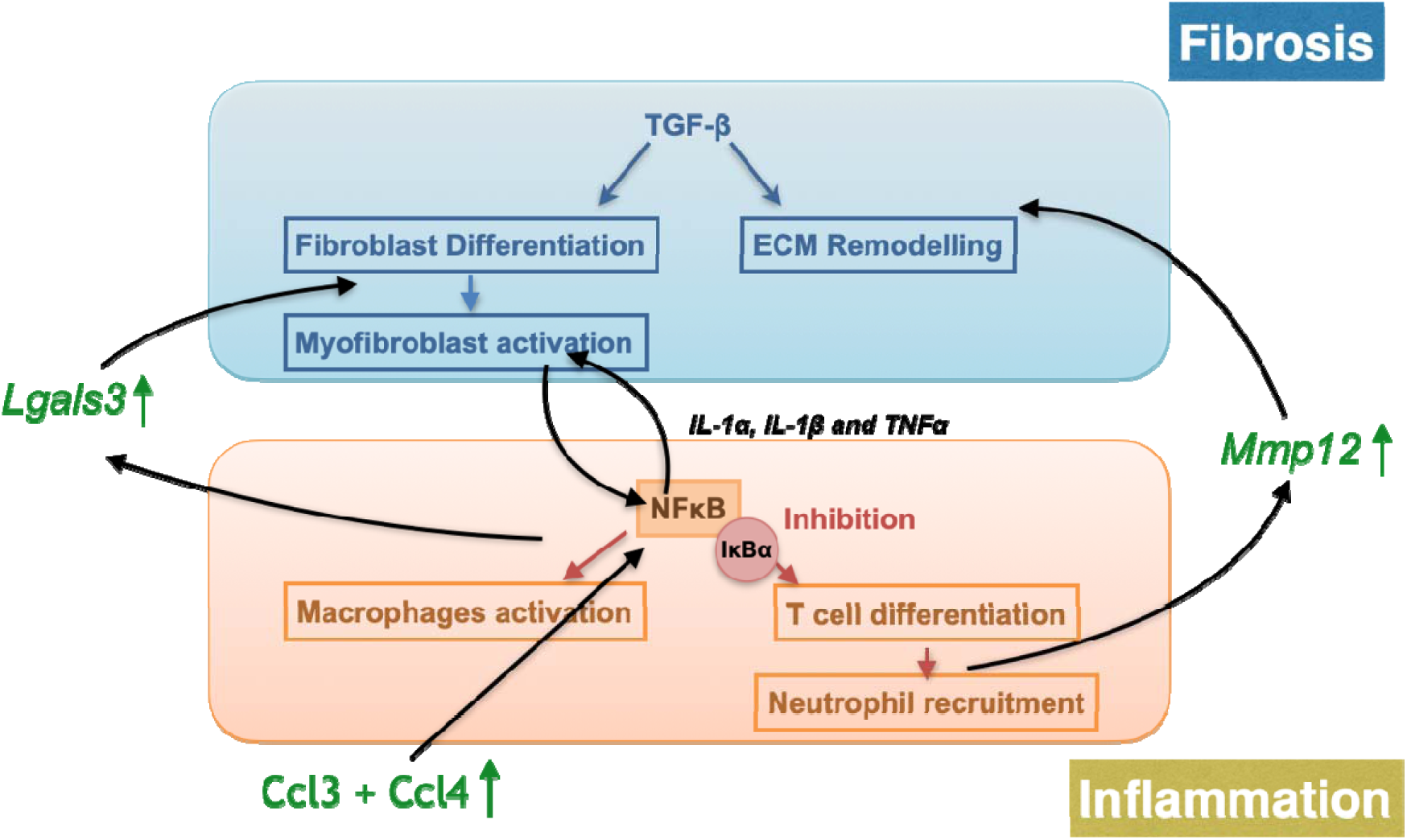
Proposed NPC cardiac fibrosis mechanism related to NFkB-mediated inflammation in cardiomyocytes. Summary chart of proposed mechanism on how transcriptomic data implicate molecular pathways linking fibrosis and inflammation, two key biological processes that may contribute to disease progression in Niemann-Pick type C (NPC). **Top panel** depicts the proposed fibrosis pathway, centerd on Transforming Growth Factor Beta (TGF-β), which promotes fibroblast differentiation, myofibroblast activation, and extracellular matrix (ECM) remodeling. These processes are associated with fibrotic remodeling. *Lgals3*, a gene known to modulate TGF-β signaling and fibroblast activation, is upregulated in NPC hearts. **The bottom panel** outlines the inflammation pathway, focused on NF-κB signaling, which is normally inhibited by IκBα. Upon release, NF-κB activates macrophages, promotes T cell differentiation and drives neutrophil recruitment. *Ccl3* and *Ccl4* chemokines involved in immune cell recruitment, are also upregulated in NPC hearts. *Mmp12*, a gene involved in ECM degradation and inflammatory remodeling, is also upregulated in *Npc1^-/-^* hearts. Arrows connecting the two panels suggest potential cross-talk between inflammation and fibrosis, indicating that chronic inflammatory signaling may promote fibrotic remodeling, and vice versa.

DEGs related to cardiomyopathies, signaling pathways, ECM organisation and TGF-beta activation are plotted using String (String-db.org, Figure A8), which show protein-protein interaction and complex network relationships. From this, several genes were identified which show strong correlations indicated by the number of connecting lines. Two main clusters were formed. In Cluster 1 (Figure A8, red box), the DEGs’ functions involved are mainly DCM, HCM, ARVC, and ECM-receptor interaction. Cluster 2 DEGs (Figure A8, yellow box) are mainly related to DCM, HCM, adrenergic signaling and cardiac muscle contraction. Analysis suggests an active cardiac remodeling pathway in NPC disease at the transcriptomic level.

## Discussion

In our study a significant proportion of patients with NPC display ECG and echocardiographic abnormalities associated with conduction disease, ventricular wall thickening and impaired systolic function (all patient data can be found in Table 1, Table 2 and Appendix Table 1). Further investigations using *Npc1^-/-^* adult mice demonstrated age-related glycosphingolipid accumulation in ventricular tissue accompanied by fibrotic remodeling. ECG from Langendorff-perfused *Npc1^-/-^* mouse hearts presented QT prolongation as well as atrioventricular conduction abnormalities under isoprenaline stress (Figure 3). Transcriptomic analysis of *Npc1^-/-^* hearts revealed significant gene expression changes, suggesting a potential link between NPC disease, inflammation, fibrosis, and arrhythmogenesis.

NPC is a progressive lysosomal disorder (Patterson, 2000). The phenotypic presentation of NPC is highly variable, and ranges from a rapidly progressive neonatal form to a slowly evolving adult-onset neurogenerative condition (Las Heras *et al*., 2023). Signs and symptoms include hepatosplenomegaly, pulmonary infiltrates, neurological complications such as epileptic seizures and ataxia, psychiatric manifestations, and early-onset dementia (Las Heras *et al*., 2023). Cardiac phenotypes and symptoms have been described in multiple sphingolipidoses including Fabry disease, Gaucher disease and Niemann-Pick disease A/B (ASMD). However, to date, there have been no published reports of cardiac remodelling or events in NPC patients and therefore merits study.We began by measuring HR and identifying ECG abnormalities in a high proportion of NPC patients (8 males, 6 females). Events related to QRS complex size, fascicular block and T wave inversion were measured. Transthoracic echocardiography identified many examples of impaired LV systolic function and increase in interventricular septal thickening or LV mass which would be highly unusual in such a young patient cohort (median age of 37 [34-41] years old) and in the absence of hypertension. Physiological left ventricular hypertrophy is found in athletic individuals undergoing exercise training; however, this is unlikely to be applicable to those with a neurodegenerative disorder. Heart failure with reduced ejection fraction (HFrEF) is also uncommon at this age (Bellanca *et al*., 2023). To determine whether cardiac pathology may be NPC related rather than a comorbidity, we studied cardiac structure and function in *Npc1^-/-^* mouse hearts, a pathogenic variant found in 95 % of cases of patients diagnosed with NPC disease *Npc1^-/-^* mice presented a lifespan of approximately three months due to their more extreme phenotype. We first observed that *Npc1^-/-^* hearts (N=6, Figure 1) had and increase in sphingolipid storage consistent with lysosomal storage characteristic of NPC patients. We conducted a comprehensive structural and morphological investigation of *Npc1^-/-^* hearts and identified cardiac abnormalities, including an increased connective tissue-to-cardiac tissue ratio in the ventricles, indicative of fibrotic remodeling, as well as an elevated left atrial-to-ventricular volume ration (Figure 2). The histological evidence of enhanced cardiac fibrosis in *Npc1^-/-^* mice was further supported by transcriptomic analysis, which revealed numerous DEGs associated with cardiac fibrosis, inflammation, and activation of signaling pathways implicated in cardiac disease.

Hepatic and renal fibrotic remodeling are common features in NPC patients (Kelly *et al*., 1993; Patel *et al*., 2021). Multiple studies indicate that the degree of fibrosis in heart diseases is linked to slowed conduction and greater susceptibility to arrhythmias (de Jong *et al*., 2011). Through histological staining, fibrosis was found in *Npc1^-/-^* mouse cardiac tissue, a likely underlying cause of arrhythmogenesis. Multiple genes related to dialated cardiomyopathy (DCM) and hypertrophic cardiomyopathy (HCM) have been detected through KEGG analysis (Figure 6). HCM is a feature of Fabry disease (Seo *et al*., 2016). Furthermore, Gb3, a glycosphingolipid that accumulates in Fabry’s disease (Tuttolomondo *et al*., 2021), is also elevated in *Npc1^-/-^* cardiac mouse tissue. We also observed significant left atrial enlargement in *Npc1^-/-^* mice, although not in the echocardiographic data from NPC patients. This may be related to elevated filling pressure in response to left ventricular hypertrophy and increased LV mass, which was more marked in the mouse model but also seen in NPC patients.

Cardiac fibrotic remodeling might be triggered by several factors (Kazbanov *et al*., 2016). Our study found a high level of immune-inflammatory responses in *Npc1^-/-^* cardiac tissue, which is in line with the systematic inflammation observed in other organs and serum from NPC patients (Vanier, 2010). Taking into account the transcriptomic network analysis, the link between inflammation and fibrosis might be via TGF-β activation and NFκB-regulated inflammation. Two genes, *Mmp12* and *Lgals3*, are among the top 10 upregulated DEGs at 3, 7, 9-weeks-old, and were linked with *Tgfb1*. *Lgals3* was assessed via RT-qPCR and presented higher expression in *Npc1^-/-^* cardiac tissue compared to WT (Figure A7). *Mmp12* is activated by initial neutrophil infiltration and facilitates neutrophil clearance followed by TGF-β activation which initiates fibrosis (Mouton *et al*., 2018a). *Lgals3*, activated by M2 macrophages, has been associated with TGF-β receptor entrapment and TGF-β activation (Suthahar *et al*., 2018a). In the heart, TGF-β can increase fibrosis by activating myofibroblast differentiation which then promotes inflammation via releasing cytokines such as IL-1α, IL-1β and TNFα (Leask, 2007; Fan *et al*., 2012). These cytokines are the major activator for NFκB signaling pathway (Fiordelisi *et al*., 2019). The NFkB pathway activation can be confirmed by upregulation of the *Ccl3* and *Ccl4* genes in transcriptomic (Kim *et al*., 2017). NFκB regulates multiple inflammatory responses including macrophage activation and neutrophil recruitment, which creates a positive feedback loop for *Mmp12* and *Lgals3* activation. IκB, inhibitor of kB, is the main regulatior of the NFκB pathway. Its phosphorylation and degradation are activated by the IKK complex, which leads to activation of NFκB (Karin, 1999). While our findings suggest that inflammation may be the primary driver of cardiac fibrosis in NPC, further molecular studies are warranted to elucidate the underlying mechanisms and validate this hypothesis.

Fibrosis is a major risk factor for both brady and tachy-arrhythmias (Frangogiannis, 2021). It acts as an electrical barrier, increasing current resistance and thereby influencing cardiac conduction velocity (Kohl & Camelliti, 2012; Unudurthi *et al*., 2014). Heterogeneous fibrosis can lead to non-uniform conduction, creating a substrate for re-entrant ventricular arrhythmias (Herring *et al*., 2019). The impact of fibrosis depends on its location, fibrosis near the atrioventricular node or bundle branches can result in varying degrees of heart block, and fascicular block has been reported in some NPC patients. ECG experiments from *Npc1^-/-^* hearts revealed an increased incidence of first- and second-degree heart block in response to Iso stress (50 nM, Figure 3). The observed slowing of atrial conduction may also be linked to both atrial enlargement and fibrotic remodeling (Spach *et al*., 2007).

Prolonged QT interval was also observed in *Npc1^-/-^* hearts under β-adrenergic receptor stimulation which may reflect either slowed conduction due to fibrosis, or abnormal ion channel activity leading to increased depolarisation or reduced repolarisation. NPC was reported as the first human disease associated with defective lysosomal Ca^2+^ uptake (Lloyd-Evans *et al*., 2008; Schröder *et al*., 2010). A fall in lysosomal Ca^2+^ concentration in *Npc1^-/-^* mouse tissue was independent of lysosomal pH (Lloyd-Evans *et al*., 2008). Our transcriptomic studies confirm an up-regulation of the lysosomal H^+^-ATPase, *Atp6v0d2*. This supports the hypothesis that lower lysosomal [Ca^2+^] is unlikely the result of reduced proton-motive force. The QT prolongation observed in *Npc1^-/-^* maybe a result of ion imbalance (eg. Ca^2+^), this remains to be further explored in detailed single cell electrophysiological studies. NAADP is a downstream mediator of β-adrenergic activation which facilitates the release of Ca^2+^ from lysosomes and enhances excitation-contraction coupling and cardiac contraction (Capel *et al*., 2015). Conditions of reduced lysosomal Ca^2+^ may therefore lead to impaired myocardial contractility and/or prolongation of the QT interval (Meng et al., 2023). Indeed, impaired systolic function was observed in two NPC patients. Prolongation of QT interval is associated with increased risk of polymorphic ventricular tachycardia, particular torsades de pointes, as well as sudden cardiac death (Herring et al., 2019).

The survival of patients with NPC at all ages has improved by approximately ten years due to miglustat therapy (Patterson *et al*., 2020). Cardiac complications, particularly fibrosis and arrhythmias, are progressive conditions that often take years to manifest. In this study, we observed increased glycolipid accumulation and upregulation of inflammation and cardiac-related genes with increasing age. This may explain why cardiac symptoms have historically been overlooked in NPC patients. With effective treatments such as extending lifespan and mitigating neurological symptoms, the underlying cardiac pathological remodeling becomes increasingly unmasked as a significant long-term risk.

Miglustat has been shown to effectively control the neurological symtoms of NPC patients (Patterson *et al*., 2020). However, depite this therapeutic benfit, some NPC patients still experience sudden death events (Walterfang *et al*., 2012; Zhao *et al*., 2022), although the exact cause remains unclear. One hypothesis is that these events may be linked to increase seizure activity associated with NPC, as individuals with epilepsy have two-to three-fold higher risk of unexpected and early death compared to the general population (Asatryan, 2021). Based on our findings, another possibility is that sudden death in NPC may be triggered by malignant ventricular tachycardia or bradyarrhythmias. These events can themselves cause anoxic seizures, sometimes leading to misdiagnosis as epilepsy, or may be triggered during the postictal phase, resulting in *sudden unexpected death in epilepsy* (SUDEP) (Ha *et al*., 2025). It is worth noting that seizure activity can cause a surge in catecholamines, accompanied by hypoxia and QTc prolongation (Pang *et al*., 2025), which can also be influenced by certain antiepileptic drugs (Feldman & Gidal, 2013). These factors, when superimposed on an abnormal cardiac structural and electrophysiological substrate, as observed in both *Npc1^-/-^* adult mice and NPC patients, have the potential to promote ventricular arrhythmogenesis (Herring *et al*., 2019).

### Limitations

We acknowledge several limitations in the present study. First, the transcriptomic analyses we performed on the mouse hearts, was using bulk RNA sequencing, which captures averaged gene expression across heterogeneous cardiac tissue and does not permit definitive attribution of changes to specific cell types. To mitigate this, we integrated complementary methodologies on mouse hearts: global fibrosis assessment provided tissue-level structural context, lipid profiling offered insight into metabolic alterations beyond transcriptional changes, and *ex vivo* ECG analyses demonstrated functional electrophysiological consequences. Collectively, these multimodal data on the mouse hearts strengthen biological interpretation despite the inherent constraints of bulk transcriptomics. We also recognise that mouse *in vivo* echocardiography and telemetry would provide valuable longitudinal and physiological insight. While echocardiography was not performed on our mouse *Npc1^-/-^* model, direct functional assessment was achieved through *ex vivo* electrophysiology alongside transcriptomic and histological analyses, together supporting robust conclusions regarding cardiac remodelling and dysfunction in NPC. Telemetry, though highly informative, requires specific ethical approval, infrastructure, and funding, and will be incorporated in future studies alongside single-cell transcriptomic approaches to further refine mechanistic understanding.

### Conclusion

In conclusion, we describe for the first time, notable cardiovascular phenotypes in both NPC patients and a mouse model of the disease (knockout of *Npc1*), likely driven by activation of fibrotic pathways. These findings warrant more extensive clinical studies, as they have the potential to inform patient management and identify new therapeutic avenues for NPC treatment. A family history of cardiovascular disease including hyperlipidaemia, arrhythmias or cardiomyopathy should be carefully reviewed as part of the clinical assessment of adults with NPC. We recommend routine annual ECG screening of NPC patients, with follow-up transthoracic echocardiography and cardiac MR (magnetic resonance imaging) if ECG abnormalities are detected, and initiation of appropriate treatment until further data become available. Further clinical prospective studies on a large cohort of NPC patients is required to evaluate the findings in the context of the disease modifying therapies.

## Funding

RABB is funded by a Sir Henry Dale Wellcome Trust and Royal Society Fellowship (109371/Z/15/Z) and RABB acknowledges support from The Returning Carers’ Fund (Oxford University, Medical Sciences Division) and acknowledges research funds from the Ellis T Davies Fellowship Endowment, University of Liverpool. RABB is a Senior Research Fellow of at Linacre College. FMP is a Wellcome Trust Investigator in Science and a Royal Society Wolfson merit award holder. DAP was funded by the Mizutani Foundation. NH is a British Heart Foundation Senior Clinical Research Fellow (FS/SCRF/20/32005). DBS is a British Heart Foundation Senior Basic Science Fellow (FS/17/55/33100 and FS/SBSRF/22/31022). DBS acknowledges support from Additional Ventures, the John Fell Oxford University Press Research Fund, and the Oxford British Heart Foundation Centre of Research Excellence Core Infrastructure Fund (RE/18/3/34214). ABO acknowledges support from UK Research and Innovation grant No. 10110728. RB was funded by Wellcome Trust (218514/Z/19/Z), Merck Sharp and Dohme Corp. and Janssen Pharmaceutica NV. For the purpose of open access, the author has applied a CC BY public copyright licence to any Author Accepted Manuscript version arising from this submission.

## Ethical Considerations

This investigation conforms to the principles outlined in the Declaration of Helsinki. This study was registered as a service evaluation within our NHS Trust. It involved a retrospective analysis of data collected during routine clinical care. As such, it did not require ethical approval, which was confirmed by the Trust’s Research and Innovation Department. No patients were recruited prospectively for the purposes of this study. Patients diagnosed with NPC under the care of the Adult Inherited Metabolic Diseases Department in Salford underwent 12-lead ECG and transthoracic echocardiography as part of a registered health improvement project at Salford Royal Hospital (Registration number: 25HIP12).

## Conflict of Interests

The authors declare no competing interests.

## Acknowledgements

We thank the Oxford Centre for Histopathology Research for help with histological services. We also thank Dr Zhaozheng Meng for support with histology image segmentation in the early stages of this project. Abstract figure was prepared using FigureLab under a licensed paid account held by the authors and edited in powerpoint.

## Author Contributions

RABB and FMP conceived the research. RABB, FMP raised research funding for the study, RAC raised funds for RNA Seq, RABB and FMP supervised all experimentation and study design. RAC, QS and IP conducted electrophysiology experimentation. QS conducted all lipids studies and omics analysis. EA helped with RT qPCR analysis. QS carried out animal dissections and cell isolations. DBS, LB and VSR conducted histology experiments. NH and ML contributed to mouse ECG experiments. RABB, QS, and IP wrote the first draft. DP, CS, DS contributed to mouse breeding and sample collection studies. KMS, NH, PW and RS conducted human clinical research. AB-O wrote code for histology segmentation studies and ran analysis. TA and QS conducted molecular studies. RB conducted additional histology and qPCR experiments. All authors have contributed to the content and refinement of the manuscript. All authors contributed intellectually to the study.

## Cover Graphic

Niemann-Pick Disease Type C (NPC) is associated with cardiac fibrosis and inflammation. Patient findings showed a high prevalence of ECG abnormalities. NPC mouse heart studies showed glycosphingolipid accumulation, fibrosis (TGF-β, myofibroblast activation, Lgals3, Mmp12) and inflammation (NF-κB, recruited macrophages/neutrophils, Ccl3/4), cardiac structural remodelling, conduction abnormalities and arrhythmogenesis. Created in FigureLabs and powerpoint.

## Appendix

**Appendix Table 1:**
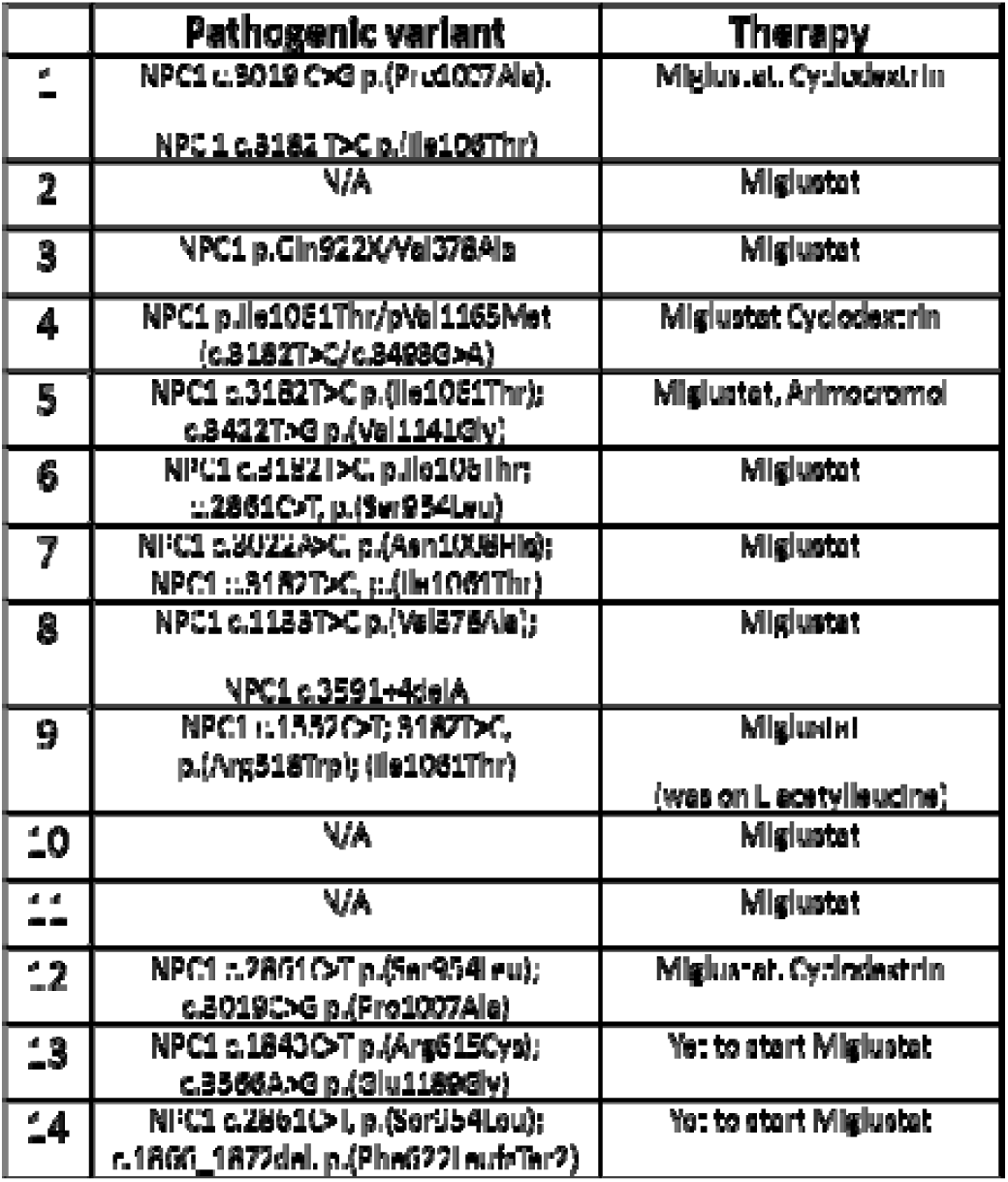
Genotypes and therapy of NPC patients. Summary table of genotype and therapy for each 14 NPC patients.

## Appendix Figure Legends

**Figure A1 related to Figure 1.**
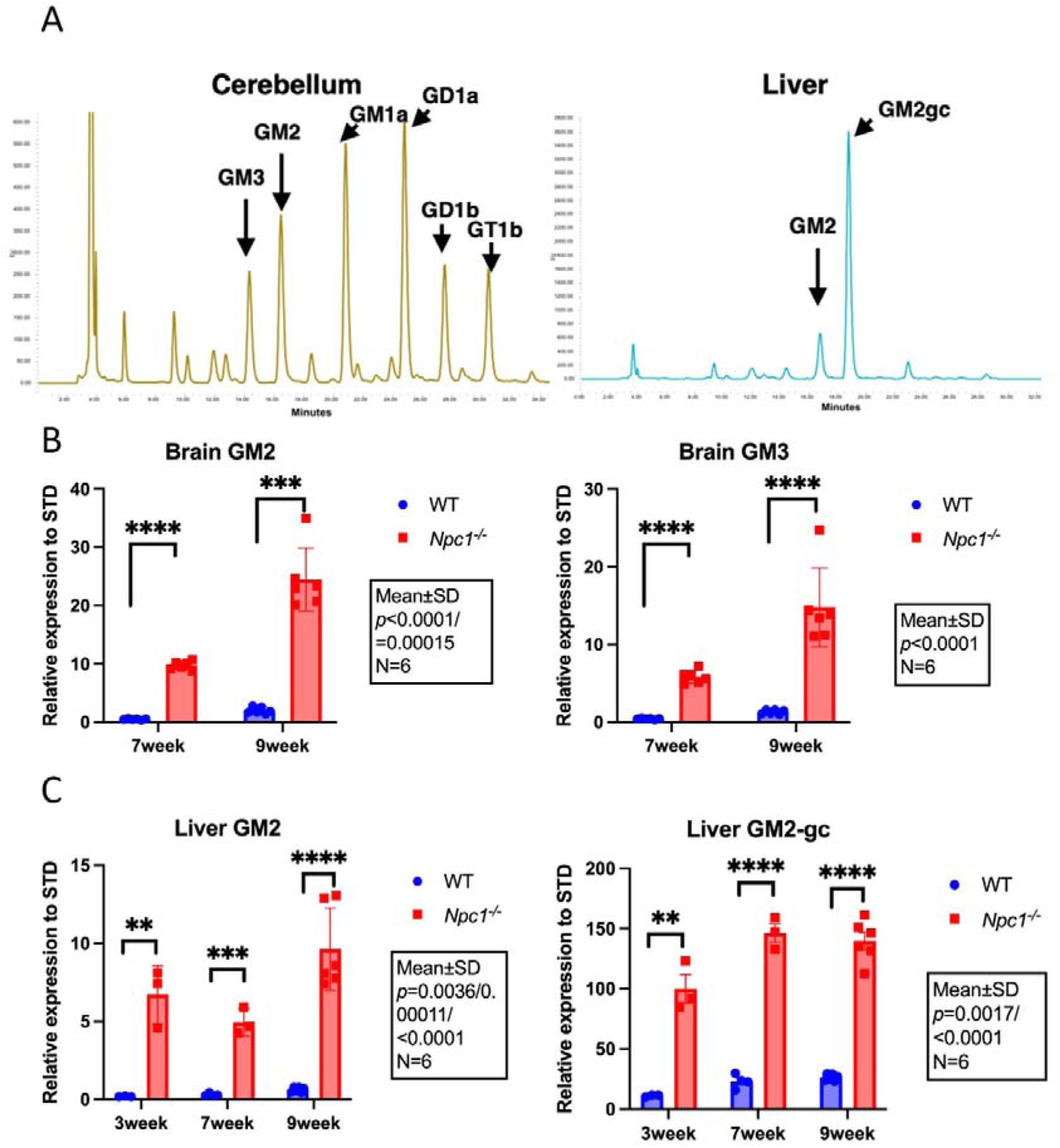
A. Representative high performance liquid chromatography traces from brain and liver of wildtype (WT, black) and *Npc1^-/-^* (red) mice. Levels of lipids in brain (B) GM2 and GM3 in WT (N=6, blue) and *Npc1^-/-^* (N=6, red) at 7 and 9 weeks and in Liver (C) at 3, 7 and 9 weeks. Data are presented as mean±SD. Student t-test. Expression level of each lipid has been normalized to standard (STD).

**Figure A2 related to Figure 2.**
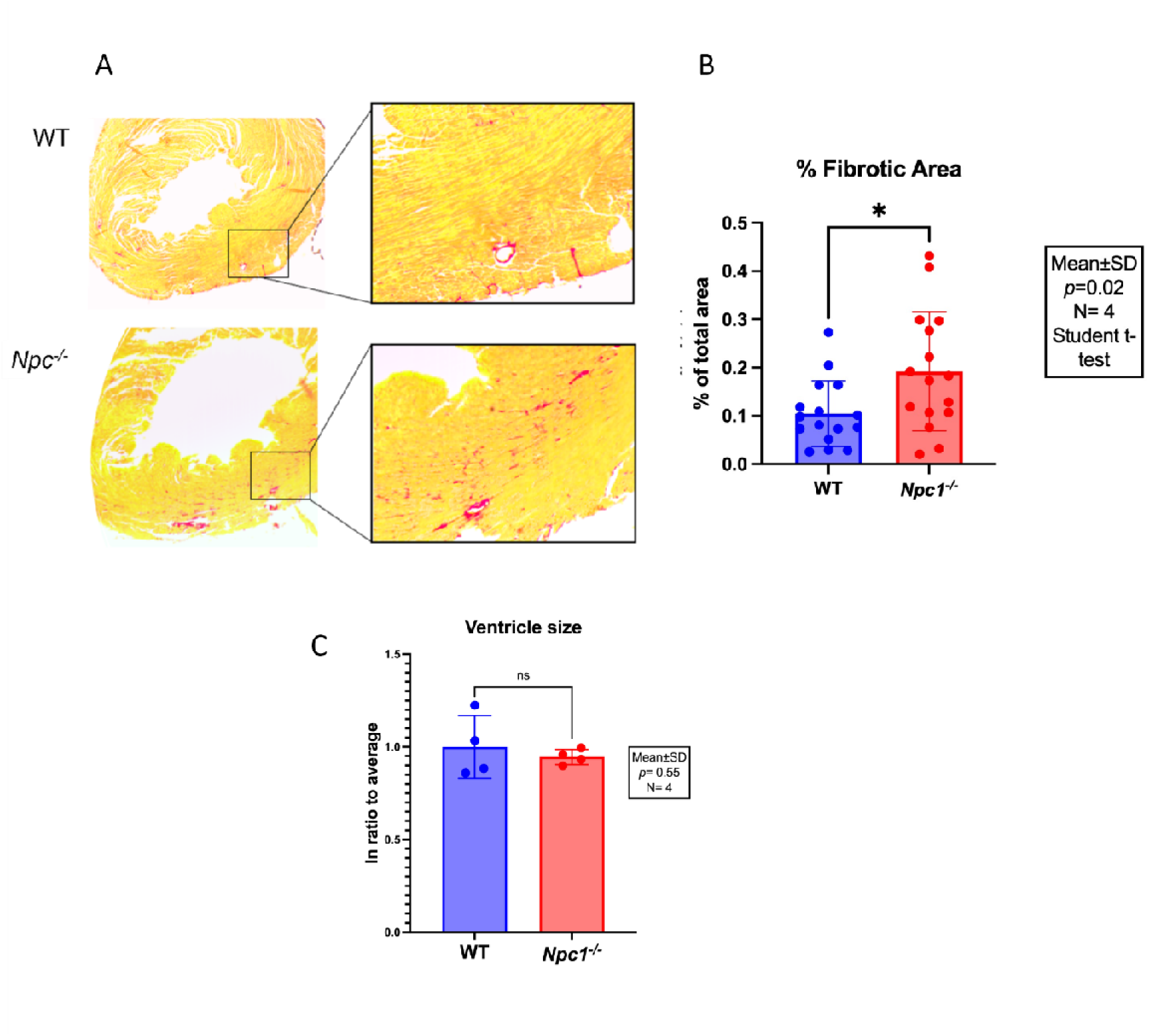
Histological studies indicates fibrosis in *Npc1^-/-^* hearts at 9-weeks. A. Picrosirius red staining of 9-week-old wildtype (WT) and *Npc1^-/-^* hearts. (A) Representative coronal sections from WT and *Npc1^-/-^* hearts. Increased frequency of collagen signal (red) not associated with vasculature or peri/epicardium are demonstrated in *Npc1^-/-^* than WT sections. B. Collagen signal determined to be consistent with fibrosis was quantified in sections from *Npc1^-/-^* (N=4) and WT (N=4) with at least four sections per heart. Analysis showed significant increase in fibrotic area (N=4, \**p*=0.02, student t-test). C. *Npc1^-/-^* vs WT hearts at 7-weeks-old. Ventricle size average of WT (N=4) *Npc1^-/-^* (N=4) mouse hearts at 7-week-old. *p*>0.05, Student t-test.

**Figure A3 related to Figure 4.**
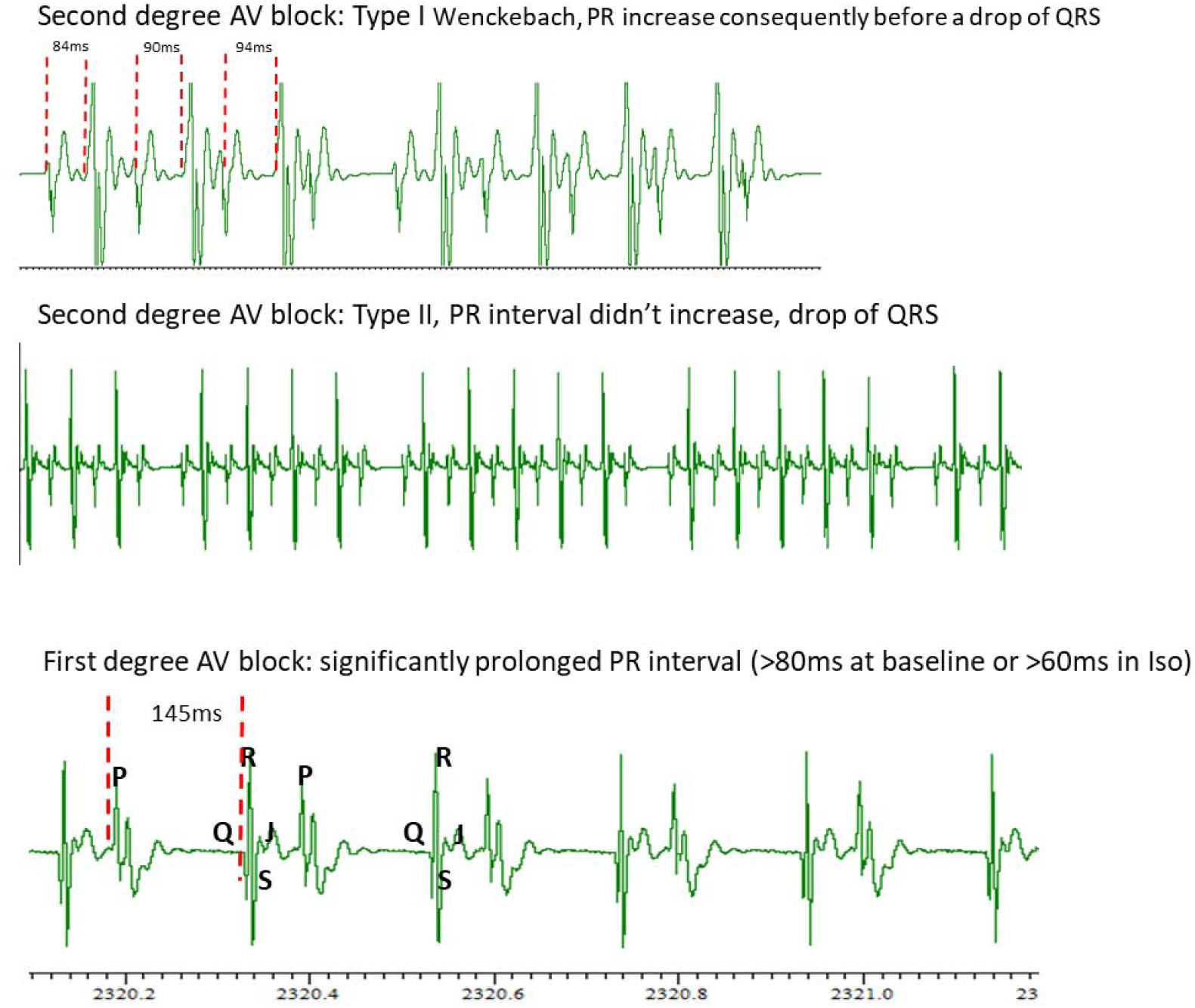
ECG trace examples of different forms of AV blocks observed in *Npc1^-/-^* hearts. (i) Second degree AV block: Type I Wenckebach, PR increase consequently before a drop of QRS. (ii) Second degree AV block: Type II, PR interval didn’t increase, drop of QRS. (iii) First degree AV block: significantly prolonged PR interval (>80 ms at baseline or >60 ms in Iso). AV: Atrioventricular.

**Figure A4 related to Figure 4.**
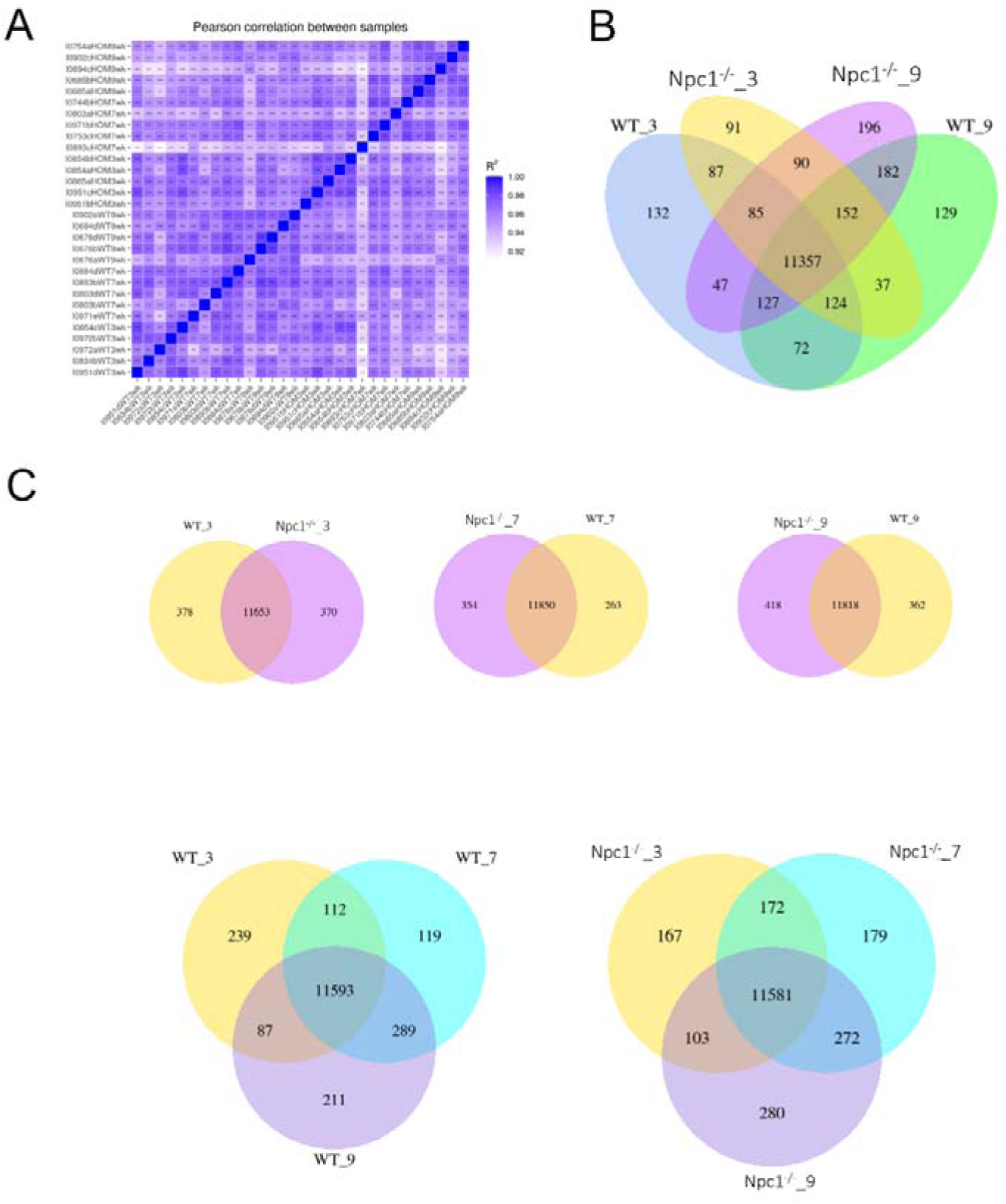
A. Pearson correlation analysis between of each sample; Gene dataset showed profile difference between each mouse. B. Co-expression analysis showing using Venn-diagram between wildtype (WT) at 3 weeks, *Npc1^-/-^*at 3 weeks, WT at 9 weeks and *Npc1^-/-^*at 9 weeks. C. Co-expression analysis showing using Venn-diagram between (i) WT and *Npc1^-/-^*at 3 weeks, (ii) WT and *Npc1^-/-^*at 7 weeks (iii) WT and *Npc1^-/-^*at 9 weeks (iv) WT at 3, 7 and 9 weeks and (v) *Npc1^-/-^*at 3, 7 and 9 weeks.

**Figure A5 related to Figure 5.**
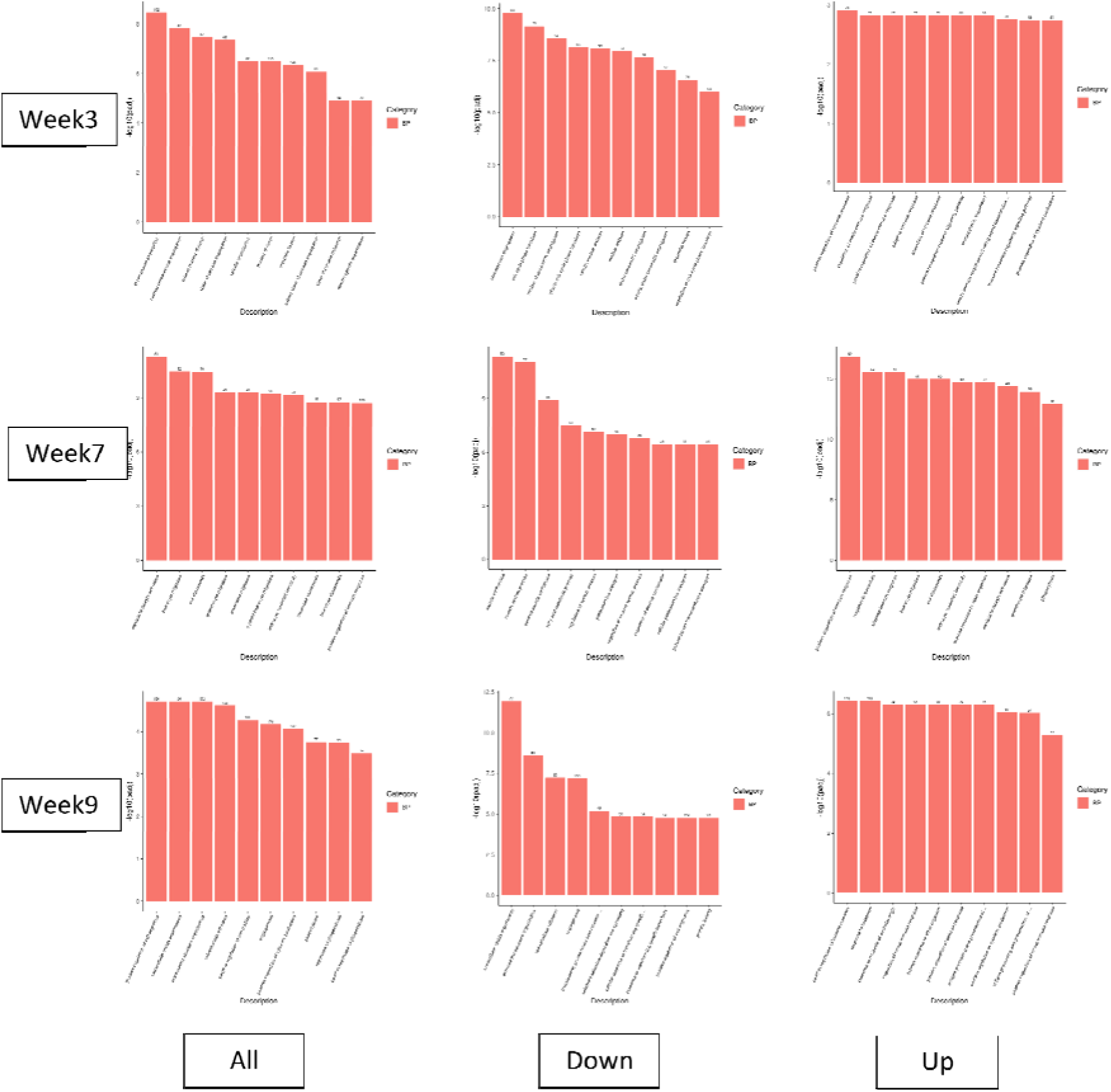
Top 10 enriched GO biological processes for DEGs in *Npc1^-/-^* compared to wildtype (WT) hearts in all DEGs, downregulated DEGs, and upregulated DEGs at 3 weeks; at 7 weeks and at 9 weeks.

**Figure A6 related to Figure 6.**
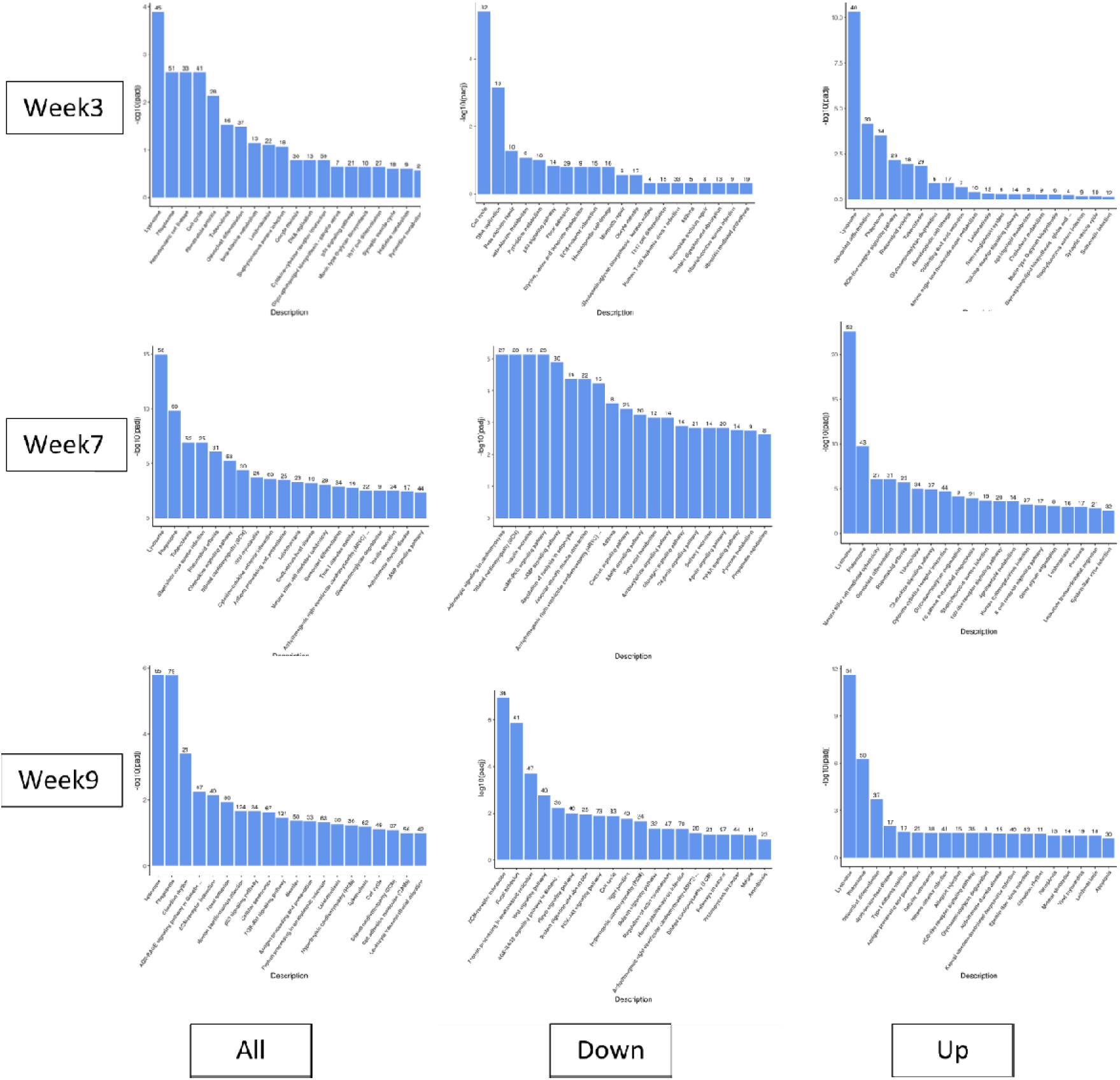
Top 10 enriched KEGG pathways for DEGs in *Npc1^-/-^* compared to wildtype (WT) hearts in all all DEGs, downregulated DEGs, and upregulated DEGs at 3 weeks; at 7 weeks and at 9 weeks.

**Figure A7 related to Figure 4.**
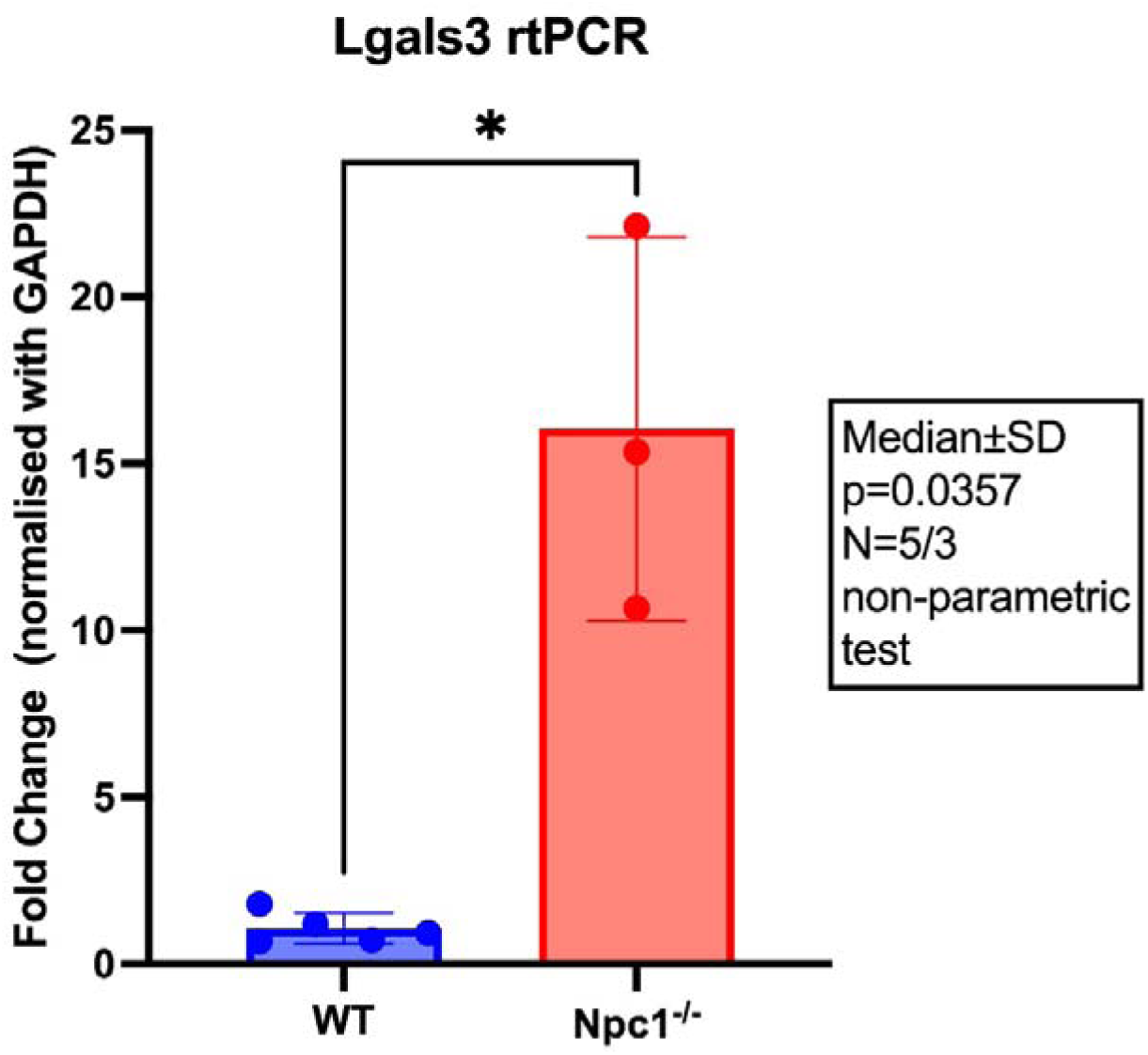
Relative expression of *Lgals3* normalized to *Gapdh* in wildtype (WT, N=5) and *Npc1^-/-^* (N=3) whole heart samples. Data are shown as fold change relative to WT. Bars represent median±SD. Statistical significance was assessed using non-parametric test;\**p*<0.05).

**Figure A8 related to Figure 4.**
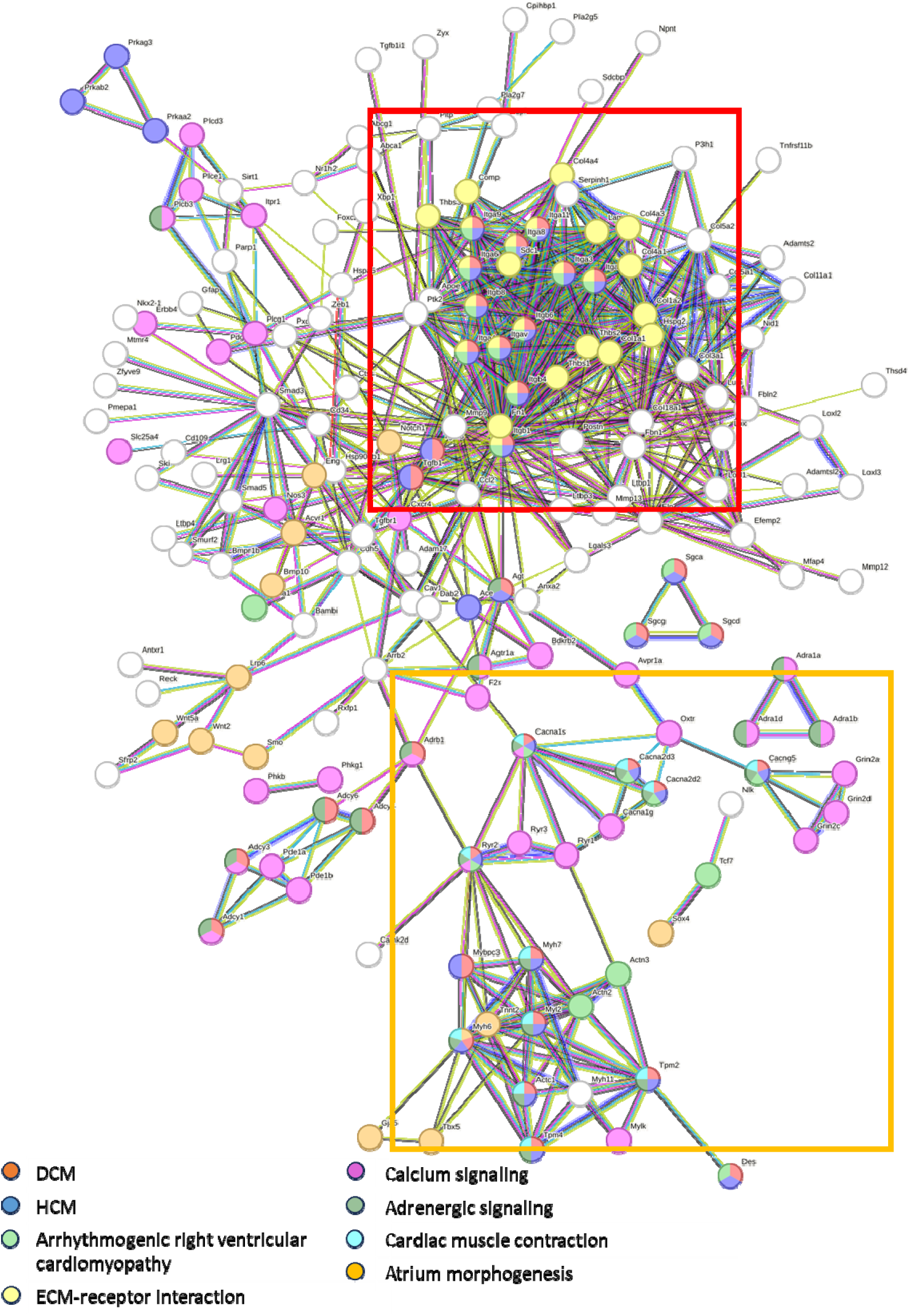
STRING network analysis on protein-protein interaction in *Npc1^-/-^* and WT hearts. Using DEGs detected by KEGG analysis in relation to cardiac disease and signaling. STRING network visualises the transcript DEGs’ protein-protein interaction. Colour of the dot represents the pathways/disease the protein is involved in. DEGs group in one cluster show close interaction with each other and therefore, point out a significant regulated pathway in NPC transcriptomic. Cluster one (red box), protein highly related to cardiomyopathies and ECM remodeling, point-out the fibrosis related gene network. Cluster two (yellow box), related to cardiac signaling pathway and muscle contraction.

## Notes

https://figshare.com/s/552a96811a6cc3991d1f

### Competing Interest Statement

The authors have declared no competing interest.

