## Appendix Table and Figures for "Molecular and Structural Basis of Cardiac Remodelling in Niemann-Pick Type C"

### Slide 1
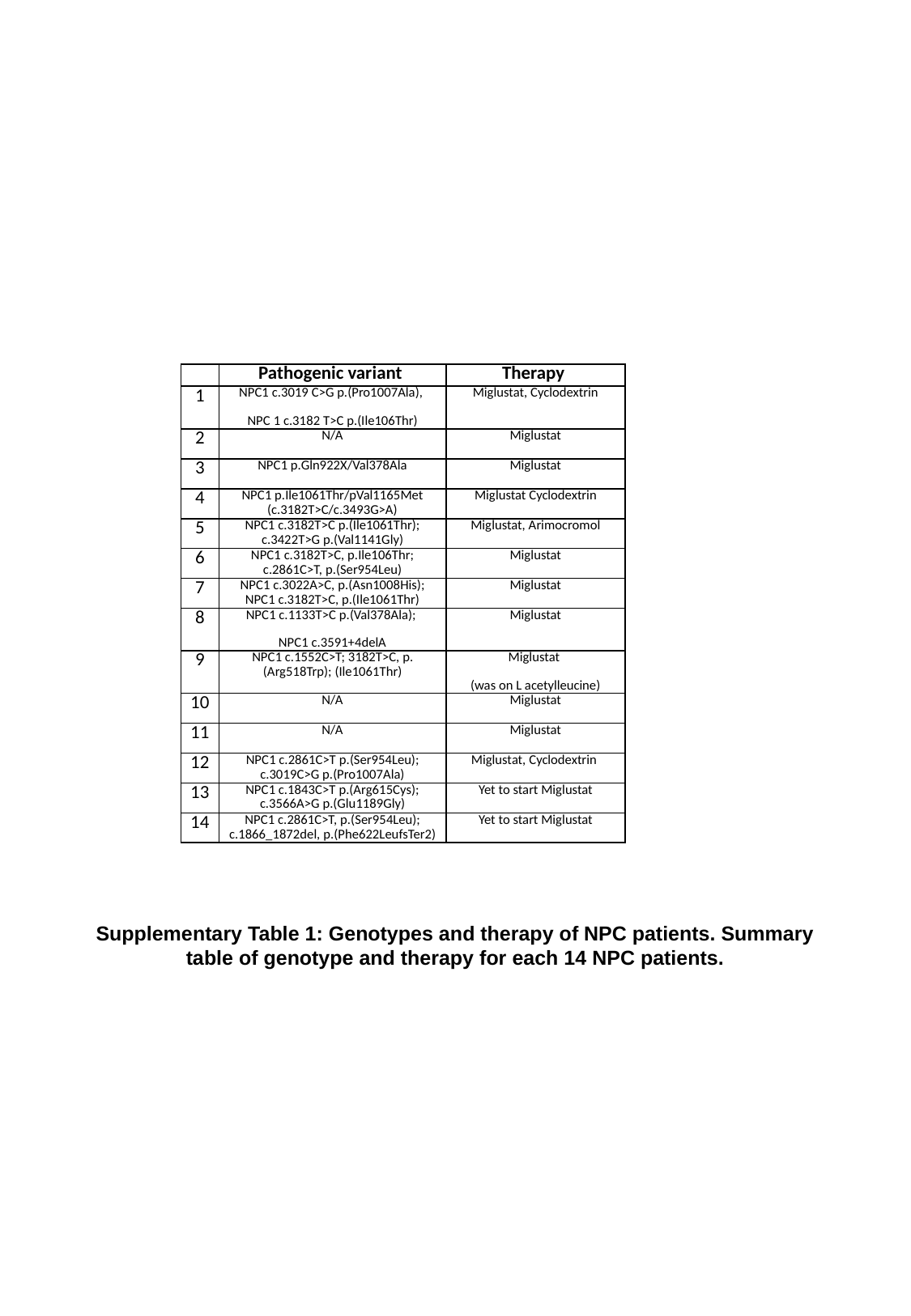

| | Pathogenic variant | Therapy |
| --- | --- | --- |
| 1 | NPC1 c.3019 C>G p.(Pro1007Ala), NPC 1 c.3182 T>C p.(Ile106Thr) | Miglustat, Cyclodextrin |
| 2 | N/A | Miglustat |
| 3 | NPC1 p.Gln922X/Val378Ala | Miglustat |
| 4 | NPC1 p.Ile1061Thr/pVal1165Met (c.3182T>C/c.3493G>A) | Miglustat Cyclodextrin |
| 5 | NPC1 c.3182T>C p.(Ile1061Thr); c.3422T>G p.(Val1141Gly) | Miglustat, Arimocromol |
| 6 | NPC1 c.3182T>C, p.Ile106Thr; c.2861C>T, p.(Ser954Leu) | Miglustat |
| 7 | NPC1 c.3022A>C, p.(Asn1008His); NPC1 c.3182T>C, p.(Ile1061Thr) | Miglustat |
| 8 | NPC1 c.1133T>C p.(Val378Ala); NPC1 c.3591+4delA | Miglustat |
| 9 | NPC1 c.1552C>T; 3182T>C, p.(Arg518Trp); (Ile1061Thr) | Miglustat (was on L acetylleucine) |
| 10 | N/A | Miglustat |
| 11 | N/A | Miglustat |
| 12 | NPC1 c.2861C>T p.(Ser954Leu); c.3019C>G p.(Pro1007Ala) | Miglustat, Cyclodextrin |
| 13 | NPC1 c.1843C>T p.(Arg615Cys); c.3566A>G p.(Glu1189Gly) | Yet to start Miglustat |
| 14 | NPC1 c.2861C>T, p.(Ser954Leu); c.1866\_1872del, p.(Phe622LeufsTer2) | Yet to start Miglustat |
Supplementary Table 1: Genotypes and therapy of NPC patients. Summary table of genotype and therapy for each 14 NPC patients.

### Slide 2
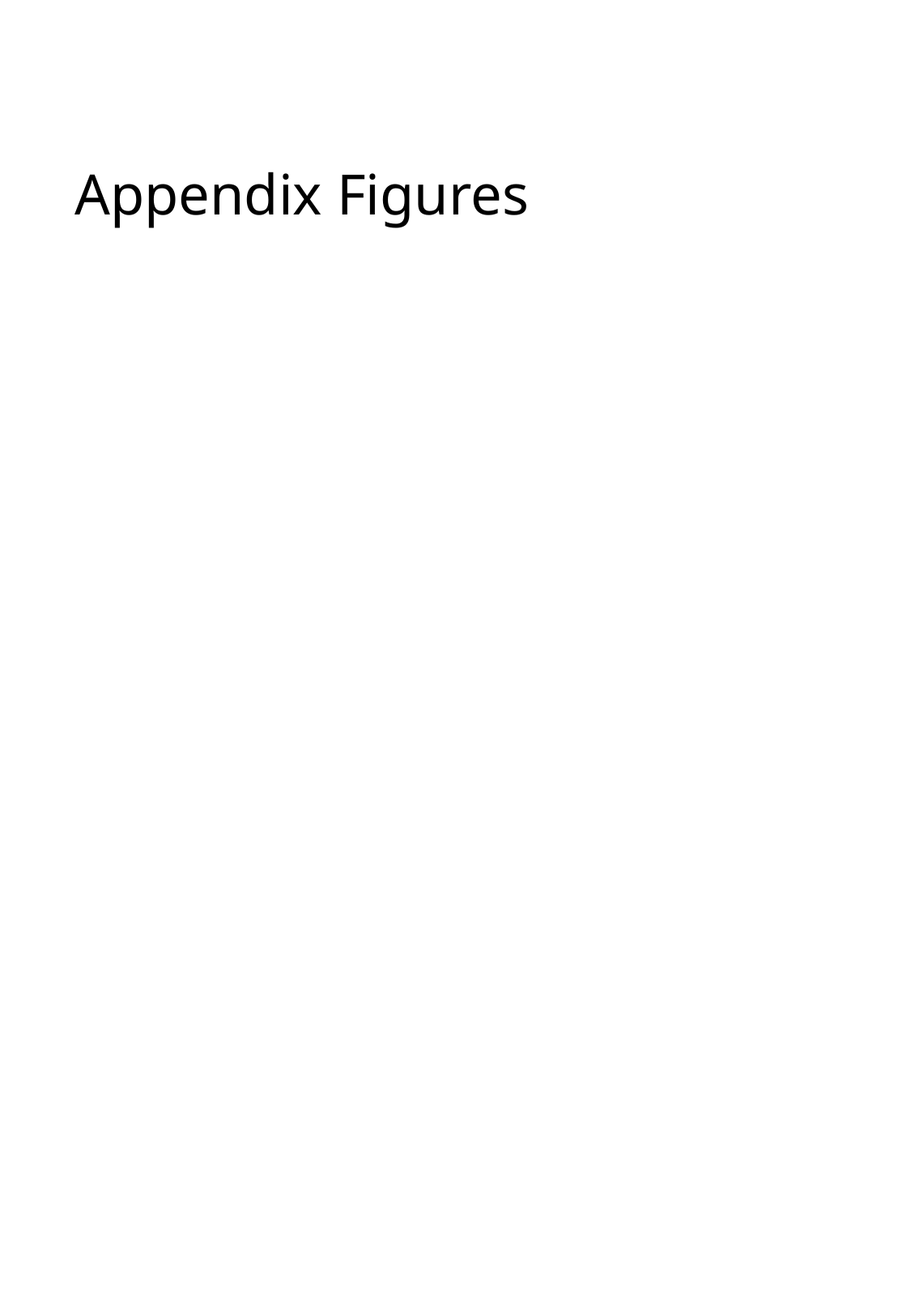

#### Appendix Figures

### Slide 3
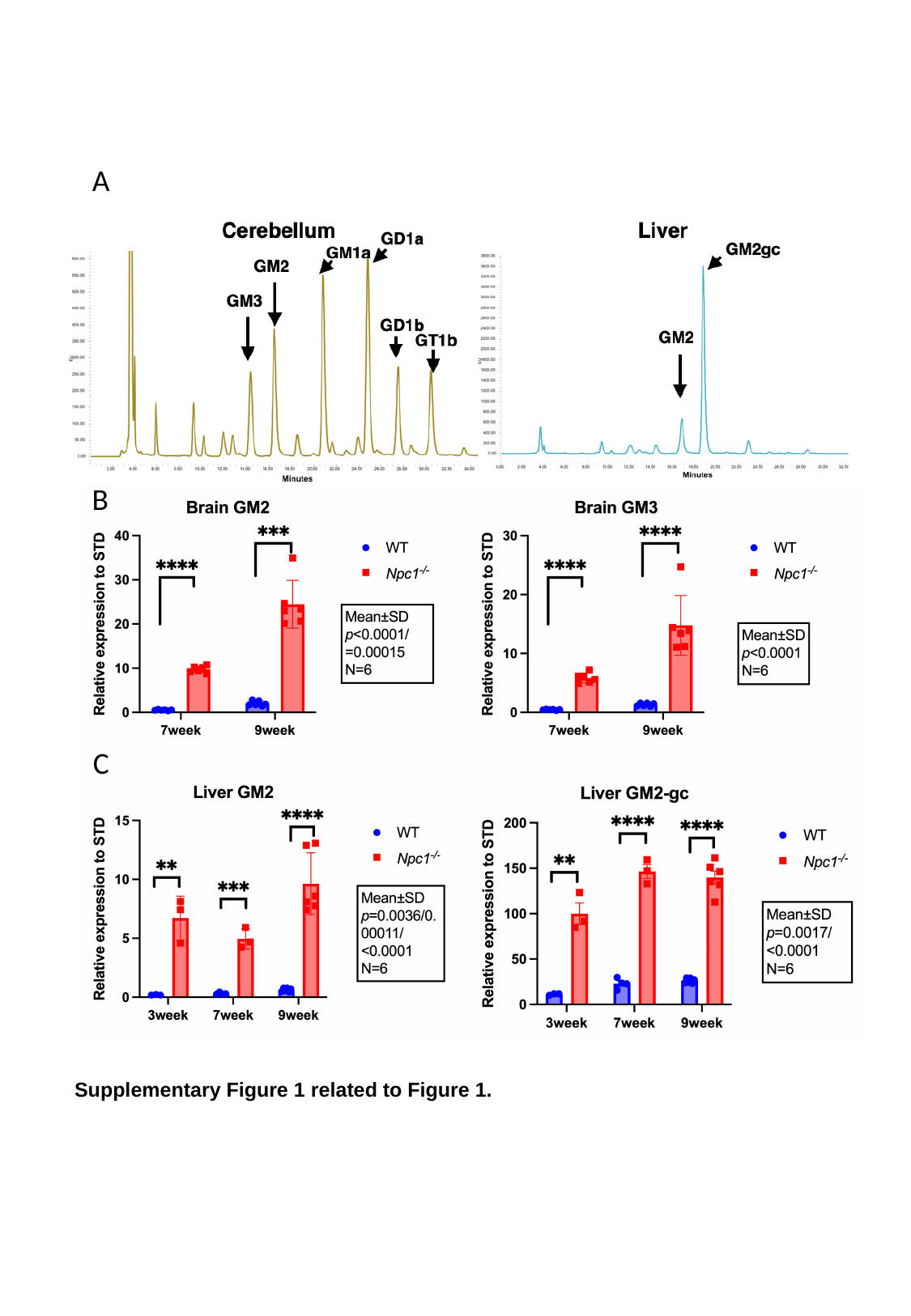

A
B
C
Supplementary Figure 1 related to Figure 1.

### Slide 4
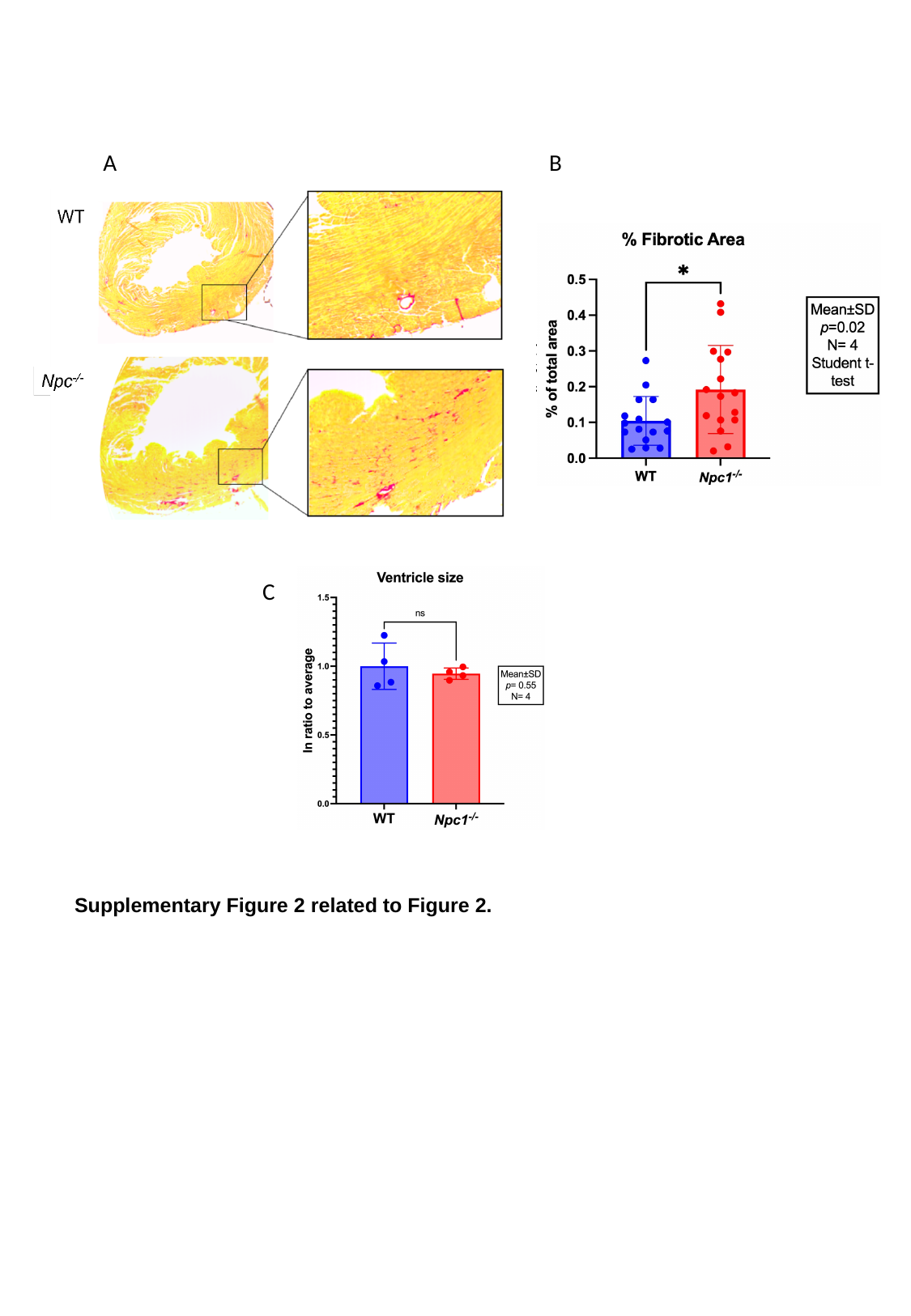

B
A
C
Supplementary Figure 2 related to Figure 2.

### Slide 5
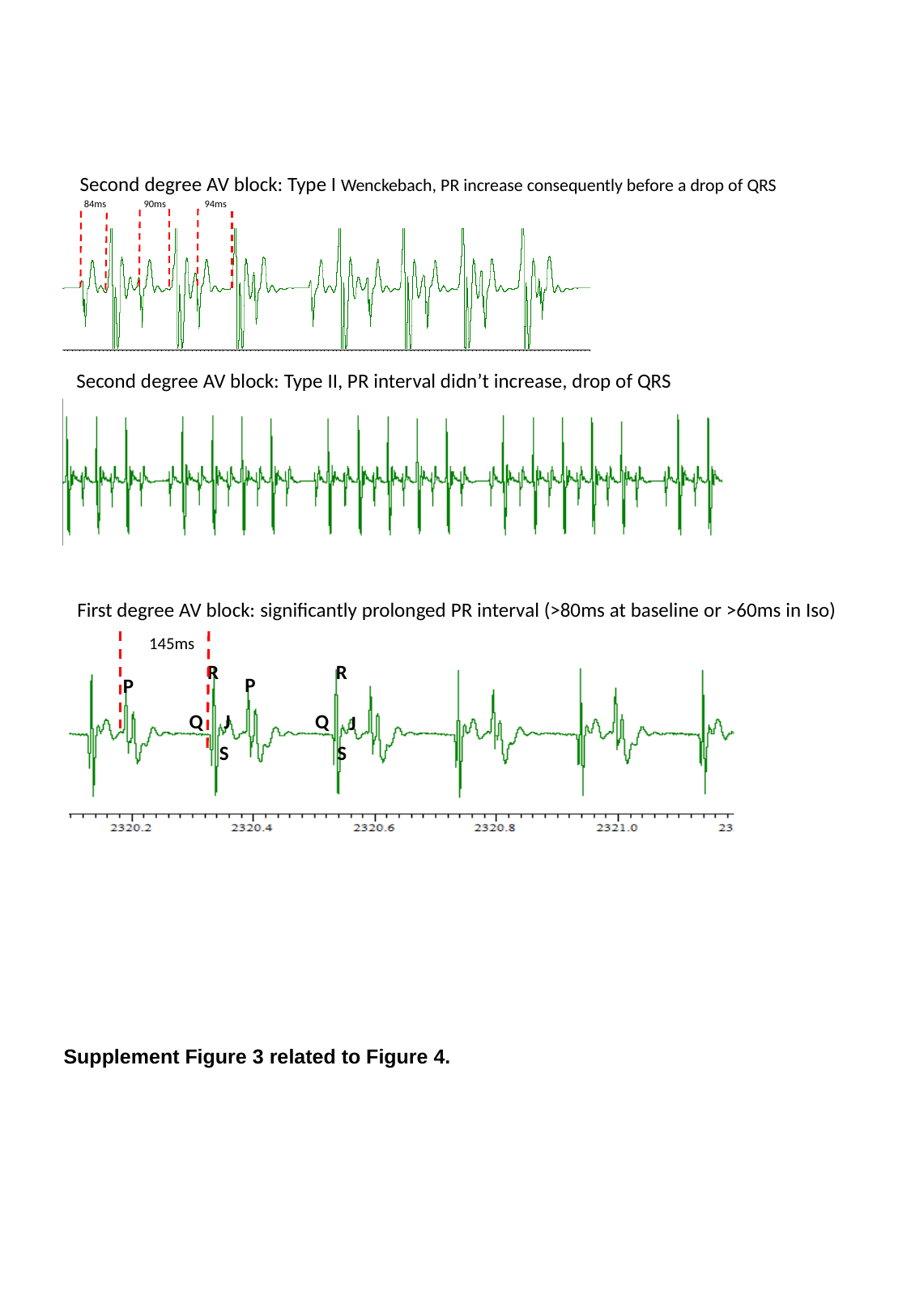

Second degree AV block: Type I Wenckebach, PR increase consequently before a drop of QRS
84ms
90ms
94ms
Second degree AV block: Type II, PR interval didn’t increase, drop of QRS
First degree AV block: significantly prolonged PR interval (>80ms at baseline or >60ms in Iso)
145ms
R
R
P
P
Q
Q
J
J
S
S
Supplement Figure 3 related to Figure 4.

### Slide 6
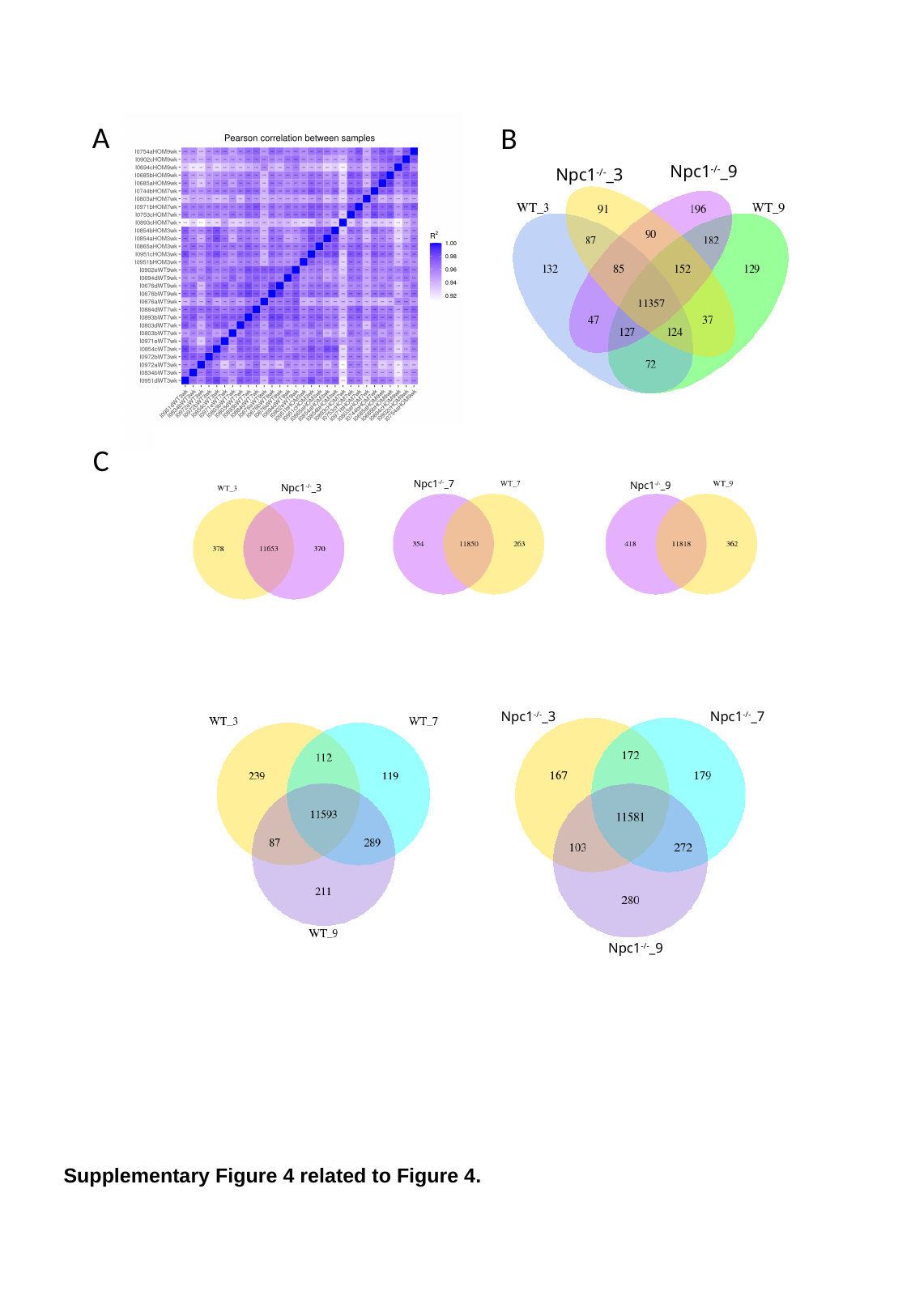

Npc1^-/-^_9
Npc1^-/-^_3
B
A
Npc1^-/-^_3
C
Npc1^-/-^_7
Npc1^-/-^_9
Npc1^-/-^_7
Npc1^-/-^_3
Npc1^-/-^_9
B
Supplementary Figure 4 related to Figure 4.

### Slide 7
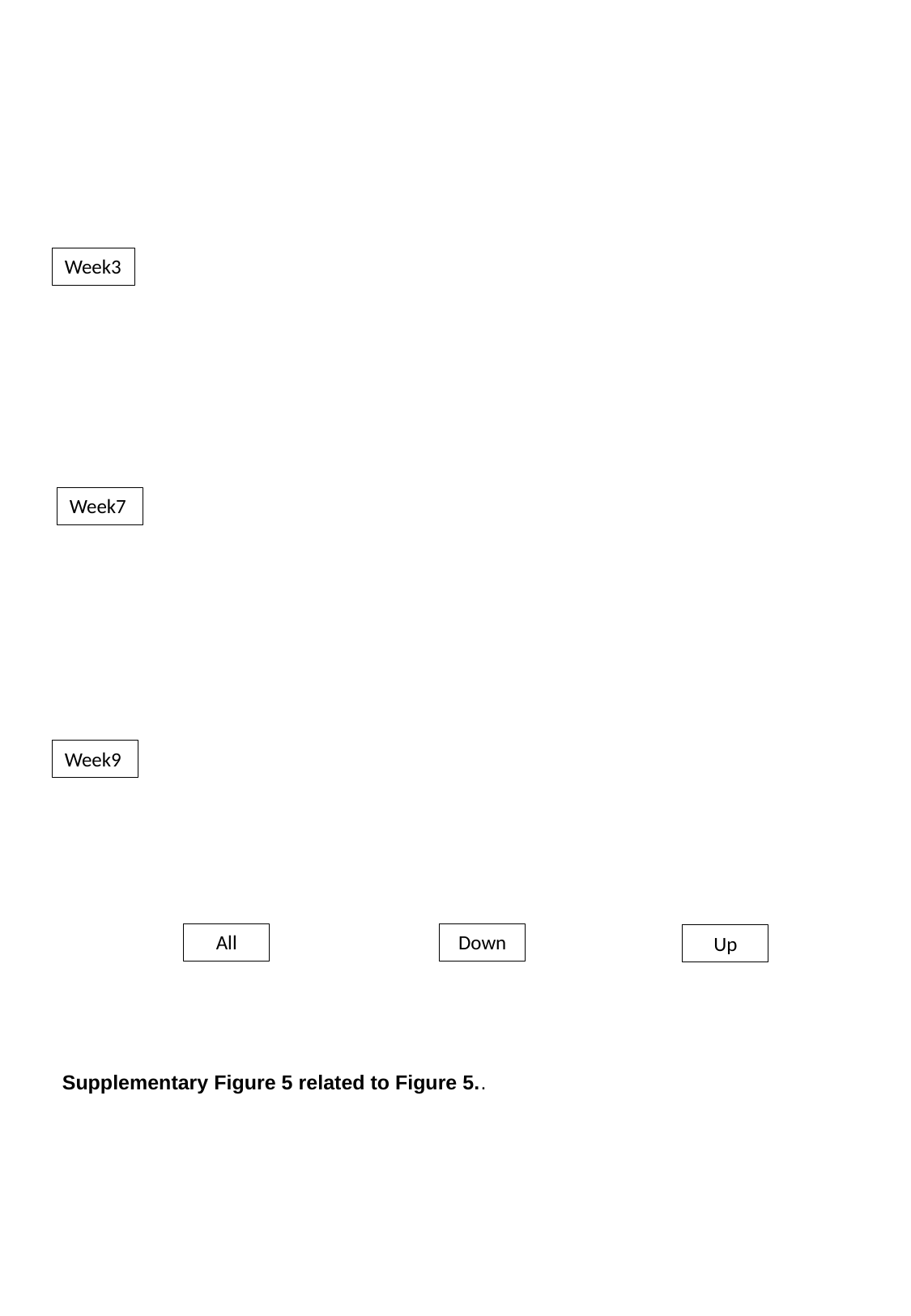

Week3
Week7
Week9
All
Down
Up
Supplementary Figure 5 related to Figure 5..

### Slide 8
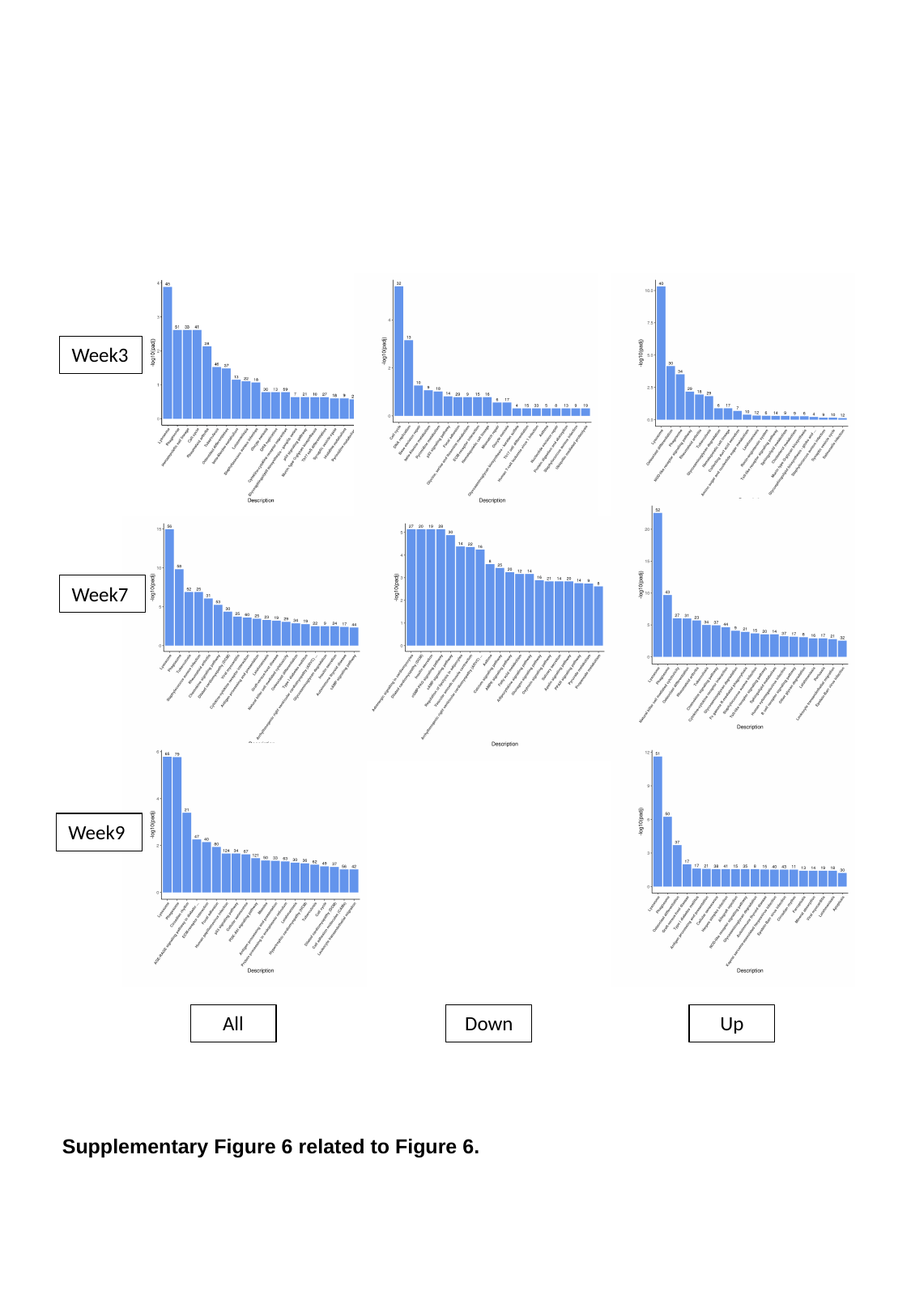

Week3
Week7
Week9
All
Down
Up
Supplementary Figure 6 related to Figure 6.

### Slide 9
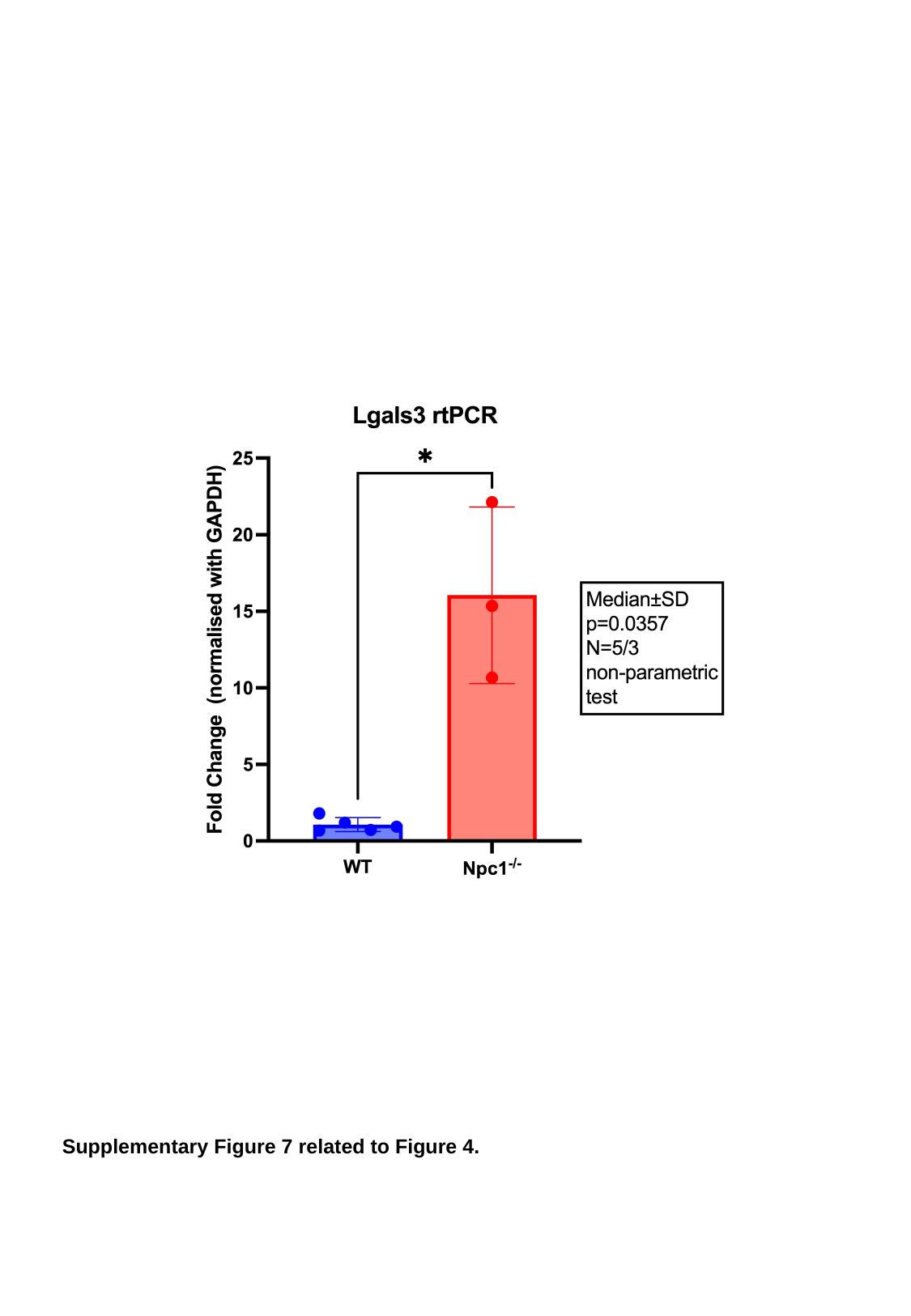

Supplementary Figure 7 related to Figure 4.

### Slide 10
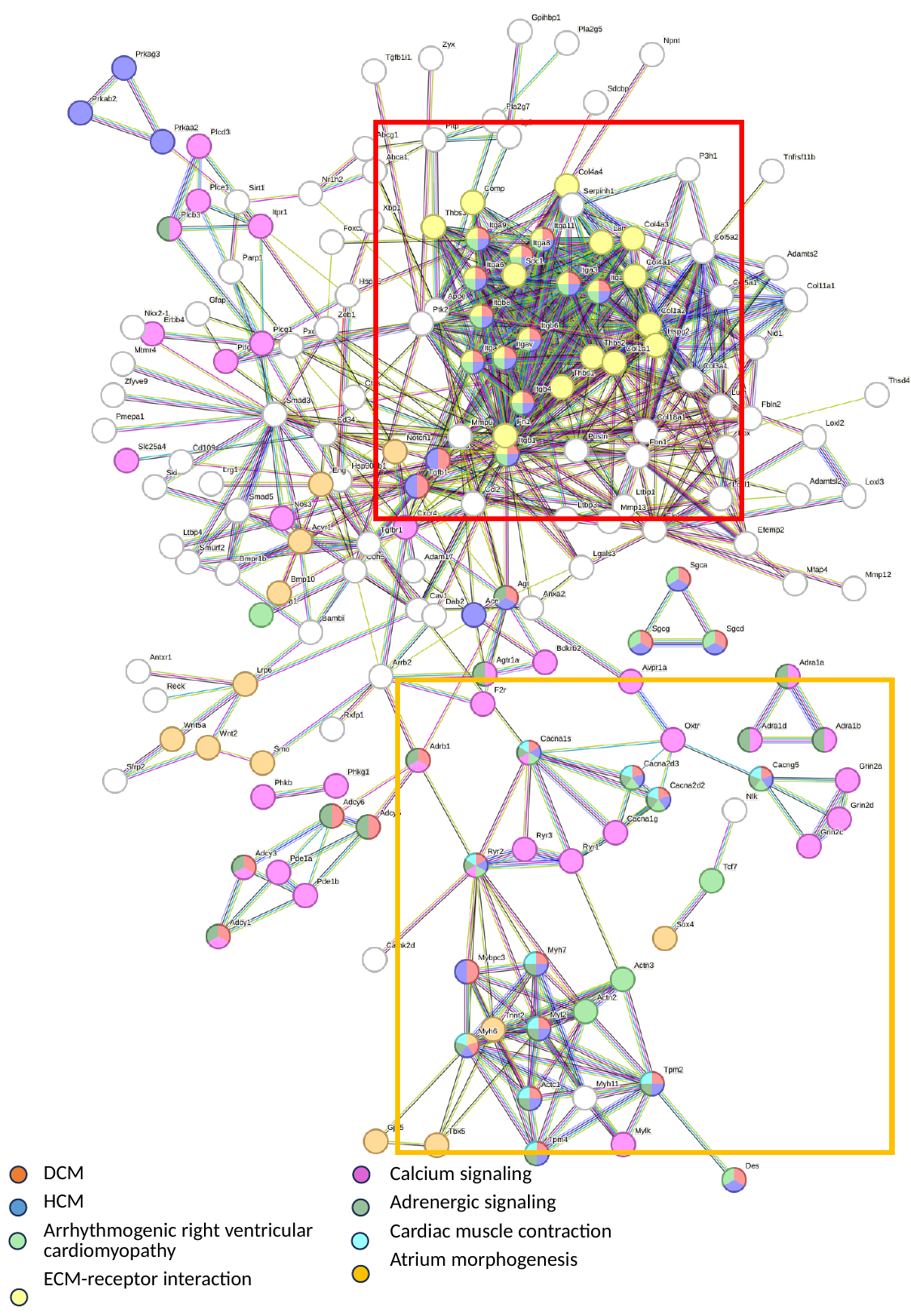

| DCM |
| --- |
| HCM |
| Arrhythmogenic right ventricular cardiomyopathy |
| ECM-receptor interaction |
| Calcium signaling |
| --- |
| Adrenergic signaling |
| Cardiac muscle contraction |
| Atrium morphogenesis |
Supplementary Figure 8 related to Figure 4.

### Slide 11
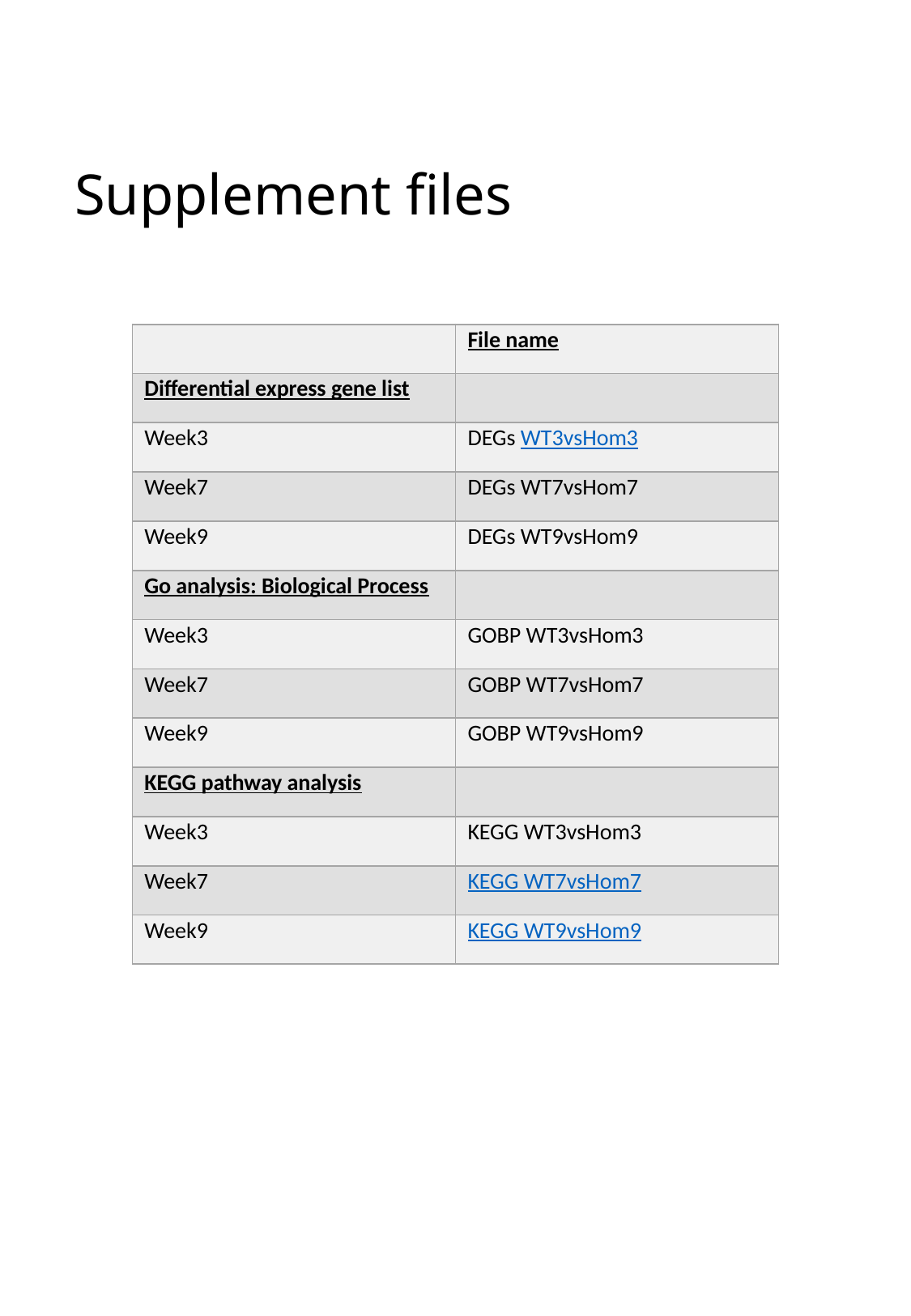

#### Supplement files
| | File name |
| --- | --- |
| Differential express gene list | |
| Week3 | DEGs WT3vsHom3 |
| Week7 | DEGs WT7vsHom7 |
| Week9 | DEGs WT9vsHom9 |
| Go analysis: Biological Process | |
| Week3 | GOBP WT3vsHom3 |
| Week7 | GOBP WT7vsHom7 |
| Week9 | GOBP WT9vsHom9 |
| KEGG pathway analysis | |
| Week3 | KEGG WT3vsHom3 |
| Week7 | KEGG WT7vsHom7 |
| Week9 | KEGG WT9vsHom9 |
